# Distinct threat and valence signals in rat nucleus accumbens core

**DOI:** 10.1101/2021.05.27.446063

**Authors:** Madelyn H. Ray, Mahsa Moaddab, Michael A. McDannald

## Abstract

Appropriate responding to threat and reward is essential to survival. The nucleus accumbens core (NAcc) is known to support and organize reward behavior. More recently our laboratory has shown the NAcc is necessary to discriminate cues for threat and safety. To directly reveal NAcc threat responding, we recorded single-unit activity from 7 female rats undergoing Pavlovian fear discrimination. Rats fully discriminated cues for danger, uncertainty, and safety. Demonstrating direct threat responding, most NAcc neurons showed greatest firing changes to danger and uncertainty. Heterogeneity in cue and reward firing led to the detection of multiple, functional populations. One NAcc population specifically decreased firing to threat (danger and uncertainty). A separate population bi-directionally signaled valence through firing decreases to negative valence events (danger and uncertainty) and opposing firing increases to positive valence events (reward and safety onset). The findings point to the NAcc as a neural source of threat information and a more general valence hub.

## Introduction

Appropriate responding to reward and threat is essential to survival. Rather than leave encounters with these positively and negatively valenced events to chance, animals form predictions about their occurrence. The nucleus accumbens plays a well-established role in learning about and organizing behaviors towards positively valenced outcomes and their predictive cues, particularly for food rewards (Baldo and Kelley, 2007; Carlezon and Thomas, 2009; Basar et al., 2010; Mannella et al., 2013; Klawonn and Malenka, 2018). The nucleus accumbens is primarily comprised of two functionally and anatomically distinct subregions: the core and the shell (Voorn et al., 1989; Zahm and Brog, 1992). The nucleus accumbens core (NAcc) is central to reward-related behavior. NAcc lesions or pharmacological manipulations decrease responding to reward-predictive cues (Parkinson et al., 1999a, 2000; Di Ciano et al., 2008; Floresco et al., 2008; Blaiss and Janak, 2009; Chaudhri et al., 2010; Ambroggi et al., 2011; Corbit and Balleine, 2011; Fraser and Janak, 2017). NAcc neurons encode various aspects of reward (Roesch et al., 2009; Krause et al., 2010; McGinty et al., 2013; Sugam et al., 2014; Sicre et al., 2020). For example, NAcc neurons show greater firing increases to a reward-predictive cue compared to a neutral cue (Cerri et al., 2014), and respond to a discriminative stimulus that signals reward availability (Ambroggi et al., 2011). In addition to signaling palatability (Roitman et al., 2005; Taha, 2005), NAcc neurons track relative reward value across a variety of conditions (Ottenheimer et al., 2018).

However, the NAcc is not exclusively involved in positive valence and is necessary for negatively valenced learning and behavior. Discrimination procedures utilizing sucrose and quinine outcomes have found that a substantial portion of nucleus accumbens neurons acquire firing to quinine and its predictive cue (Setlow et al., 2003; Roitman et al., 2005). Accumbens lesions centered on the NAcc disrupt the increased latencies of ‘go’ responses to quinine cues (Schoenbaum and Setlow, 2003). Further, differing accumbens optogenetic simulation patterns can induce approach versus avoidance behavior (Soares-Cunha et al., 2020). However, negatively valenced tastes such as quinine and their predictive cues produce a highly specific suite of behavioral and orofacial responses (Grill and Norgren, 1978; Kerfoot et al., 2007; Berridge, 2019). Whereas threat cues associated with nociceptive stimuli, such as foot shock, produce a distinct suite of defensive behaviors: freezing, hyperventilation, piloerection and more (Bolles, 1970; Bolles and Collier, 1976; Bouton and Bolles, 1980). It remains unclear how individual NAcc neurons differentially respond to cues constituting the threat domain of negative valence.

The NAcc has not been viewed as a major node in the brain’s threat network. This title is often bestowed on the amygdala (LeDoux et al., 1990; Campeau and Davis, 1995; Killcross et al., 1997; Koo et al., 2004). While the amygdala is undoubtedly important, threat learning and behavior is normally the product of a much larger network (Beck and Fibiger, 1995; Vetere et al., 2017). This network includes traditional ‘reward’ regions (Reynolds and Berridge, 2002; Pauli et al., 2015; Bouchet et al., 2018; Groessl et al., 2018; Cai et al., 2020; Piantadosi et al., 2020; Stephenson-Jones et al., 2020; Moaddab and McDannald, 2021; Moaddab et al., 2021). The NAcc receives a prominent, direct projection from the basolateral amygdala (Christie et al., 1987; Brog et al., 1993; Li et al., 2018). Foot-shock associated cues and contexts increase NAcc immediate early gene expression (Beck and Fibiger, 1995; Thomas et al., 2002). Indeed, there is a growing body of evidence implicating the NAcc in a variety of threat processes (Iordanova et al., 2006; Martinez et al., 2008; Badrinarayan et al., 2012; Budygin et al., 2012; Wenzel et al., 2015; Zhang et al., 2020). However, studies aimed at determining the role of NAcc in fear using traditional cued and contextual conditioning procedures have produced conflicting results (Parkinson et al., 1999b; Levita et al., 2002; Schwienbacher et al., 2004; Wendler et al., 2014).

Recently, we reported an essential role for the NAcc in scaling suppression of rewarded nose poking to degree of threat (Ray et al., 2020). Using a discrimination procedure consisting of cues predicting unique foot shock probabilities: danger (*p* = 1.00), uncertainty (*p* = 0.25), and safety (*p* = 0.00), we showed the NAcc is necessary to acquire discriminative fear. We further revealed a specific role for the NAcc in the acquisition and expression of rapid uncertainty-safety discrimination. Despite these findings, it is unclear whether NAcc neurons respond to threat cues. To reveal NAcc threat responding, we recorded single-unit activity from female rats undergoing fear discrimination. The goal of the experiment was threefold. First, we sought to reveal whether NAcc neurons are generally responsive to threat cues. If so, we could then determine if NAcc threat responding patterns correspond to functions identified via NAcc lesion and optogenetic inhibition (Ray et al., 2020). Specifically, NAcc neurons sustaining activity across cue presentation would be expected to fully discriminate cues, while showing preferential firing to danger. NAcc neurons showing transient activity to cue onset would be expected to fully discriminate uncertainty and safety. Lastly, using suppression of rewarded nose poking as our dependent measure allowed us to compare threat and reward-related firing. The complete behavior/recording approach allowed us to uncover signals specifically reflecting threat or more generally reflecting valence that spans threat and reward.

## Methods

The recording/fear discrimination approach is based on prior work from our laboratory (Moaddab et al., 2021).

### Subjects

Seven female Long Evans rats were bred in the Boston College Animal Care Facility. Behavioral testing took place in adulthood when rats weighed 215-300 g. Rats were single-housed on a 12 h light/dark cycle (lights on at 6:00 a.m.) with free access to water. Rats were maintained at 85% of their free-feeding body weight with standard laboratory chow (18% Protein Rodent Diet #2018, Harlan Teklad Global Diets, Madison, WI), except during surgery and post-surgery recovery. All protocols were approved by the Boston College Animal Care and Use Committee and all experiments were carried out in accordance with the NIH guidelines regarding the care and use of rats for experimental procedures.

### Electrode assembly

Microelectrodes consisted of a drivable bundle of sixteen 25.4 µm diameter Formvar-Insulated Nichrome wires (761500, A-M Systems, Carlsborg, WA) within a 27-gauge cannula (B000FN3M7K, Amazon Supply) and two 127 µm diameter PFA-coated, annealed strength stainless-steel ground wires (791400, A-M Systems, Carlsborg, WA). All wires were electrically connected to a nano-strip Omnetics connector (A79042-001, Omnetics Connector Corp., Minneapolis, MN) on a custom 24-contact, individually routed and gold immersed circuit board (San Francisco Circuits, San Mateo, CA). The sixteen-wire bundle was integrated into a microdrive permitting advancement in ∼42 μm increments.

### Surgery

Aseptic, stereotaxic surgery was performed under isoflurane anesthesia (1-5% in oxygen). Carprofen (5 mg/kg, s.c.) and lactated ringer’s solution (10 mL, s.c.) were administered preoperatively. The skull was scoured in a crosshatch pattern with a scalpel blade to increase surface area. Six screws were installed in the skull. A 1.4 mm diameter craniotomy was performed to remove a circular skull section centered on the implant site and the underlying dura was removed to expose the cortex. Nichrome recording wires were freshly cut with surgical scissors to extend ∼2.0 mm beyond the cannula. Just before implant, current was delivered to each recording wire in a saline bath, stripping each tip of its formvar insulation. Current was supplied by a 12V lantern battery and each Omnetics connector contact was stimulated for 2 s using a lead. Machine grease was placed around the cannula and microdrive. For implantation dorsal to the NAcc, the electrode assembly was slowly advanced (∼100 μm/min) to the following coordinates relative to bregma: +1.44 mm anterior, -1.40 mm lateral, and -6.00 mm ventral from the cortex. Once in place, stripped ends of ground wires were wrapped around two screws and screws advanced to contact cortex. The microdrive base and a protective head cap were cemented in place with orthodontic resin (C 22-05-98, Pearson Dental Supply, Sylmar, CA), and the Omnetics connector was affixed to the head cap.

### Behavioral apparatus

All experiments were conducted in two behavioral chambers housed in sound-attenuating shells. Behavioral chambers had aluminum front and back walls retrofitted with clear plastic covers, clear acrylic sides and top, and a stainless-steel grid floor. Each grid floor bar was electrically connected to an aversive shock generator (Med Associates, St. Albans, VT) through a grounding device. The grounding device permitted the floor to be always grounded except during shock delivery. An external food cup and a central port, equipped with infrared photocells were present on one wall. Auditory stimuli were presented through two speakers mounted on the ceiling of enclosure. Behavior chambers were modified to allow for free movement of the electrophysiology cable during behavior; plastic funnels were epoxied to the top of the behavior chambers with the larger end facing down, and the tops of the chambers were cut to the opening of the funnel.

### Nose poke acquisition

Rats were pre-exposed to the experimental pellets in their home cages (Bio-Serv, Flemington, NJ). Rats were then shaped to nose poke for pellet delivery in the behavior chamber using a fixed ratio schedule in which one nose poke yielded one pellet. Over the next 5 days, rats were placed on variable interval (VI) schedules in which nose pokes were reinforced on average every 30 s (VI-30, day 1), and 60 s (VI-60, days 2 through 5). Nose pokes were reinforced on a VI-60 schedule throughout fear discrimination independent of cue and shock presentation.

### Fear discrimination

Prior to surgery, each rat received eight 54-minute Pavlovian fear discrimination sessions. Each session consisted of 16 trials, with a mean inter-trial interval of 3.5 min. Auditory cues were 10-s repeating motifs of patterned beep, broadband click, phaser, or trumpet (listen or download: http://mcdannaldlab.org/resources/ardbark). Each cue was associated with a unique foot shock probability (0.5 mA, 0.5 s): danger, *p* = 1.00; uncertainty, *p* = 0.25; and safety, *p* = 0.00. Auditory identity was counterbalanced across rats. For danger and uncertainty shock trials, foot shock was administered 2 s following cue offset. A single session consisted of four danger trials, two uncertainty shock trials, six uncertainty omission trials, and four safety trials; with order randomly determined by the behavioral program. After the eighth discrimination session, rats were given *ad libitum* food access and implanted with drivable microelectrode bundles. Following surgical recovery, discrimination resumed with single-unit recording. The microelectrode bundles were advanced in ∼42-84 μm steps every other day to record from new single units during the following session.

### Single-unit data acquisition

During recording sessions, a 1x amplifying headstage connected the Omnetics connector to the commutator via a shielded recording cable (Headstage: 40684-020 & Cable: 91809-017, Plexon Inc., Dallas TX). Analog neural activity was digitized, and high-pass filtered via amplifier to remove low-frequency artifacts then sent to the Ominplex D acquisition system (Plexon Inc., Dallas TX). Behavioral events (cues, shocks, nose pokes, and rewards) were controlled and recorded by a computer running Med Associates software. Timestamped events from Med Associates were sent to Ominplex D acquisition system via a dedicated interface module (DIG-716B). The result was a single file (.pl2) containing all time stamps for recording and behavior. Single units were sorted offline using principal components analysis and a template-based spike-sorting algorithm (Offline Sorter V3, Plexon Inc., Dallas TX). Timestamped spikes and events (cues, shocks, nose pokes and rewards) were extracted and analyzed with statistical routines in Matlab (Natick, MA).

### Histology

Rats were deeply anesthetized using isoflurane and final electrode coordinates were marked by passing current from a 6V battery through 4 of the 16 nichrome wires. Rats were transcardially perfused with 0.9% biological saline and 4% paraformaldehyde in a 0.2 M potassium phosphate buffered solution. Brains were extracted and post-fixed in a 10% neutral-buffered formalin solution for 24 h, stored in 10% sucrose/formalin, frozen at -80°C and sectioned via microtome. Nissl staining was performed in order to identify NAcc boundaries. Sections were mounted on coated glass slides, Nissl-stained, and coverslipped with Omnimount mounting medium (Fisher Scientific, Waltham, MA), and imaged using a light microscope (Axio Imager Z2, Zeiss, Thornwood, NY).

### Verifying electrode placement

Passing current through the wire permitted tip locations to be identified in brain sections. In addition, wire tracks leading up to tips were visible. Starting with the electrode tips, the driving path of the electrode through the brain was backwards calculated. Only single units obtained from recording locations inside of the tear-shaped region surrounding the anterior commissure (Paxinos and Watson, 2007) were considered to be in the NAcc (Figure 1C, D) and therefore only these single units were included in analyses.

**Figure 1.**
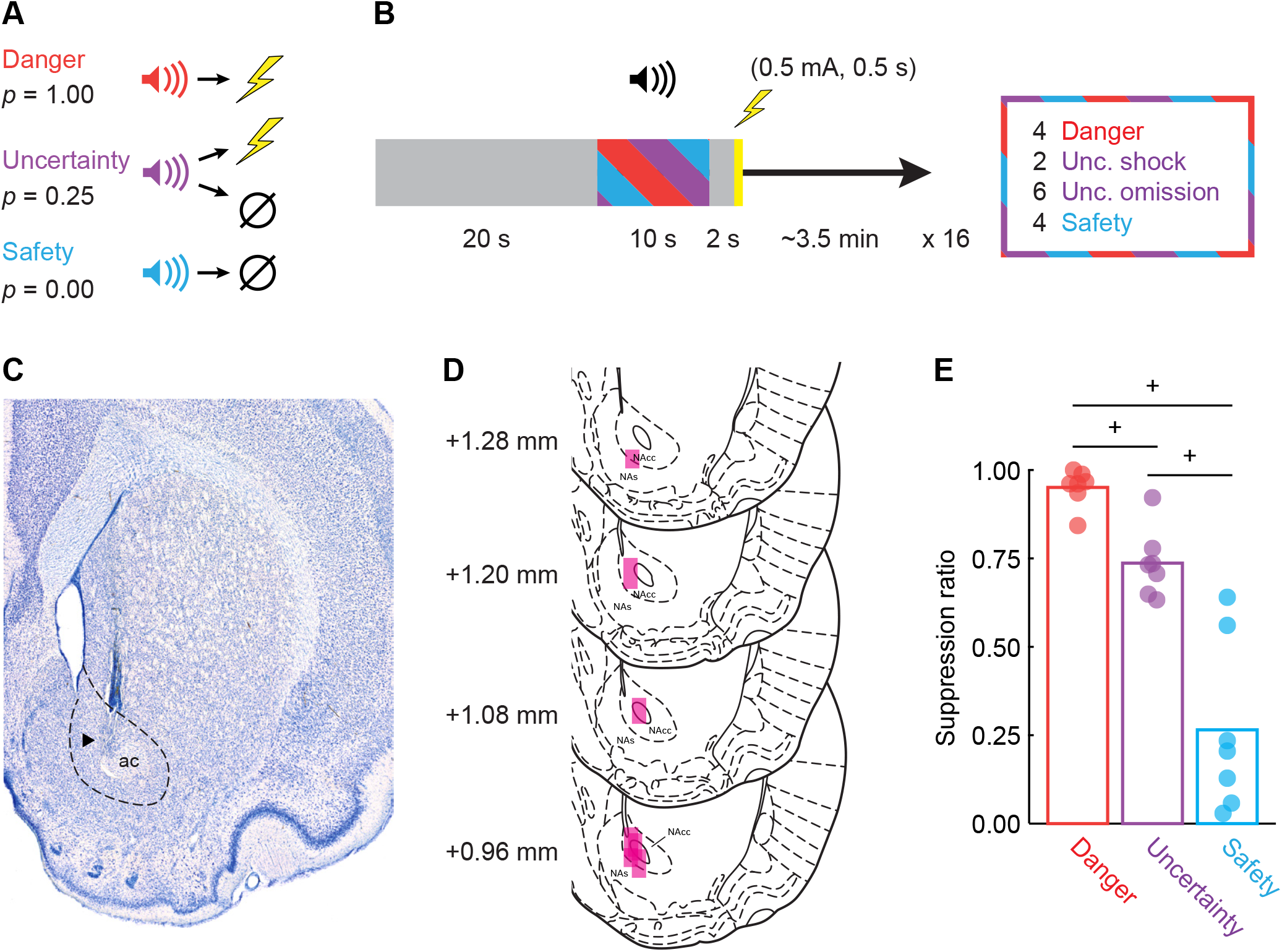
Fear discrimination, histology and behavior. (**A**) Pavlovian fear discrimination consisted of three auditory cues, each associated with a unique probability of foot shock: danger (*p* = 1.00, red), uncertainty (*p* = 0.25, purple) and safety (*p* = 0.00, blue). (**B**) Each trial started with a 20 s baseline period followed by 10 s cue period. Foot shock (0.5 mA, 0.5 s) was administered 2 s following the cue offset in shock and uncertainty shock trials. Each session consisted of 16 trials: four danger trials, two uncertainty shock trials, six uncertainty omission trials and four safety trials with an average inter-trial interval (ITI) of 3.5 min. (**C**) Example of a Nissl stained NAcc (outlined in black) section showing the location of the recording site within the boundaries of the NAcc. (**D**) Histological reconstruction of microelectrode bundle placements (n = 7) in the NAcc are represented by pink bars, bregma levels indicated. (**E**) Mean (bar) and individual subject (data points; n = 7) suppression ratio for each cue (danger, red; uncertainty, purple; safety, blue) is shown. ^+^95% bootstrap confidence interval for differential suppression ratio does not contain zero.

### 95% bootstrap confidence intervals

95% bootstrap confidence intervals were constructed for differential suppression ratios and firing, using the bootci function in Matlab. Bootstrap distributions were created by sampling the data 1,000 times with replacement. Studentized confidence intervals were constructed with the final outputs being the mean, lower bound and upper bound of the 95% bootstrap confidence interval. Null hypotheses state that differential suppression ratios (or differential firing) were not observed. Differential suppression ratios and firing – rejecting the null hypothesis – were observed when the 95% confidence interval did not include zero.

### Calculating suppression ratios

Fear was measured by suppression of rewarded nose poking, calculated as a ratio: [(baseline nose poke rate - cue nose poke rate) / (baseline nose poke rate + cue nose poke rate)]. The baseline nose poke rate was taken from the 20 s prior to cue onset and the cue poke rate from the 10 s cue period. Suppression ratios were calculated for each trial using only that trial’s baseline. A ratio of ‘1’ indicated high fear, ‘0’ low fear, and gradations between intermediate levels of fear. Suppression ratios were analyzed using analysis of variance (ANOVA) with cue (danger, uncertainty, and safety) as a factor (Figure 1E). F statistic, *p* value, partial eta squared (η_p_^2^) and observed power (op) are reported for significant main effects and interactions. Mean individual suppression ratios were visualized using the plotSpread function in Matlab (https://www.mathworks.com/matlabcentral/fileexchange/37105-plot-spread-points-beeswarm-plot).

### Identifying cue-responsive neurons

Single units were screened for cue responsiveness by comparing raw firing rate (Hz) during the 10 s baseline period just prior to cue onset to firing rate during the first 1 s and last 5 s of danger, uncertainty, and safety using a paired, two-tailed t-test (*p* < 0.05). A single unit was considered cue-responsive if it showed a significant increase or decrease in firing to any cue in either period. Bonferroni correction was not performed because this criterion was too stringent, resulting in obviously cue-responsive neurons being omitted from analysis.

### Firing and waveform characteristics

The following characteristics were determined for each cue-responsive neuron: baseline firing rate, coefficient of variance, and waveform peak-valley duration (Figure 2-3). Baseline firing rate was mean firing rate (Hz) during the 10 s baseline period just prior to cue onset. Coefficient of variance was calculated by 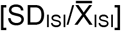, in which SD_ISI_ was the standard deviation of inter-spike interval, and 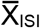 was the mean inter-spike interval. Coefficient of variance is a relative measure of the variability of spike firing, with small values indicating less variation in inter-spike intervals (more regular firing), and large values more variability (less regular firing) (Saeb-Parsy and Dyball, 2003; Moaddab et al., 2015). Waveform peak-valley duration was the x-axis distance between the valley of depolarization and the peak of after-hyperpolarization with smaller values indicating narrow waveforms (Gage et al., 2010; Gittis et al., 2011; Vachez et al., 2021).

**Figure 2.**
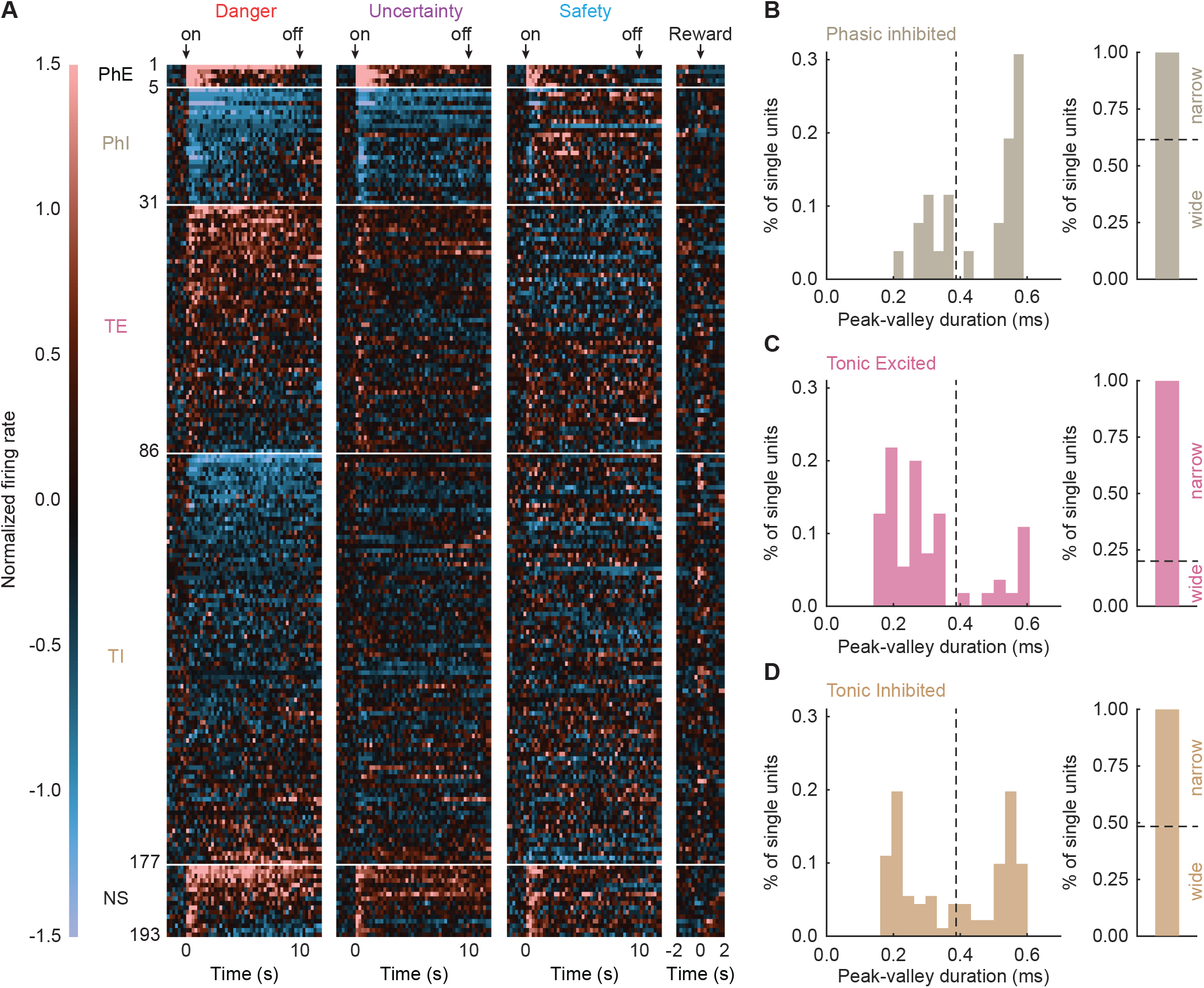
Heat plot of cue-responsive neurons. (**A**) Mean normalized firing rate for each cue-responsive neuron (n = 193) for each of the three trial types: danger, uncertainty, and safety (2 s prior to cue onset to 2 s following cue offset, in 250-ms bins), as well as reward (2 s prior to 2 s following reward delivery). Cue onset (on), offset (off), and reward are indicated by black arrows. All cue-responsive neurons are sorted by their cue-responsiveness (PhE, Phasic Excited, n = 5, black; PhI, Phasic Inhibited, n = 26, grey; TE, Tonic Excited, n = 55, pink; TI, Tonic Inhibited, n = 91, tan; NS, Non-Selective, n = 16, black). Color scale for normalized firing rate is shown to the left. A normalized firing rate of zero is indicated by the color black, with greatest increases light red and greatest decreases light blue. (**B**, left) Histogram and (**B**, right) bar graph depict distribution of the waveform peak-valley duration for Phasic Inhibited neurons. Black dashed lines divide Phasic Inhibited neurons into narrow (small values) and wide (large values) waveforms. Colors maintained from A. Identical graphs made for (**C**) Tonic Excited and (**D**) Tonic Inhibited neurons.

### K-means clustering

Clustering was performed using the Matlab kmeans function. Firing rate of all cue-responsive neurons (n = 193) was summarized in a 193 neuron x 6 epoch matrix. The 6 firing epochs were mean onset (first 1 s) and late cue (last 5 s) firing for danger, uncertainty and safety. Clustering was performed 7 times, incrementing from 1 to 7 clusters. The primary result of clustering was the cluster membership of each neuron and the mean of the squared Euclidean distance between each cluster member and the cluster centroid. Cluster number was optimized to produce the fewest number of clusters and the smallest mean Euclidean distance of each cluster member from its centroid.

### Z-score normalization

For each neuron, and for each trial type, firing rate (Hz) was calculated in 250-ms bins from 20 s prior to cue onset to 20 s following cue offset, for a total of 200 bins. Mean firing rate over the 200 bins was calculated by averaging all trials for each trial type. Mean differential firing was calculated for each of the 200 bins by subtracting mean baseline firing rate (10 s prior to cue onset), specific to that trial type, from each bin. Mean differential firing was Z-score normalized across all trial types within a single neuron, such that mean firing = 0, and standard deviation in firing = 1. Z-score normalization was applied to firing across the entirety of the recording epoch, as opposed to only the baseline period, in case neurons showed little/no baseline activity. As a result, periods of phasic, excitatory and inhibitory firing contributed to normalized mean firing rate (0). For this reason, Z-score normalized baseline activity can differ from zero. Z-score normalized firing was analyzed with ANOVA using cue, and bin as factors. F and *p* values are reported, as well as partial eta squared (η_p_^2^) and observed power (op). Reward-related firing was extracted from inter-trial intervals, when no cues were presented. Although not explicitly cued through the speaker, each reward delivery was preceded by a brief sound caused by the advance of the pellet dispenser. For reward-related firing (time locked to pellet dispenser advance), firing rate (Hz) was calculated in 250 ms-bins from 2 s prior to 2 s following advancement of pellet dispenser, for a total of 16 bins. Mean differential firing was calculated for each of the 16 bins by subtracting pre-reward firing rate (mean of 1 s prior to reward delivery).

### Heat plot and color maps

Heat plots were constructed from normalized firing rate using the imagesc function in Matlab (Figure 2, Figure 2-1). Perceptually uniform color maps were used to prevent visual distortion of the data (Crameri, 2018).

### Population and single-unit firing analyses

Population cue firing was analyzed using ANOVA with cue (danger, uncertainty, and safety) and bin (250-ms bins from 2 s prior to cue onset to cue offset) as factors (Figure 2-2, Figure 3). Uncertainty trial types were collapsed because they did not differ firing analysis. This was expected, during cue presentation rats did not know the current uncertainty trial type. F statistic, *p* value, partial eta squared (η_p_^2^) and observed power (op) are reported for main effects and interactions. The 95% bootstrap confidence intervals were reconstructed for normalized firing to each cue (compared to zero), as well as for differential firing (danger vs. uncertainty), (uncertainty vs. safety), (danger and uncertainty vs. safety), and (danger vs. uncertainty and safety) during cue onset (first 1-s cue interval) and late cue (last 5-s cue interval). The distribution of single-unit firing was visualized using a plotSpread function for Matlab. Population reward firing was analyzed using repeated measures ANOVA with bin (250-ms bins from 2 s prior to 2 s following advancement of pellet dispenser) as factor (Figure 5A-D). The 95% bootstrap confidence intervals were reconstructed for normalized firing to reward during pre (250 ms prior to reward delivery), and post (first 250 ms following reward delivery) (compared to zero), as well as for differential firing (pre vs. post). Relationships between cue firing (danger, safety) and reward (Figure 5E-G), as well as cue firing (danger vs. safety; Figure 5H-J) were determined by calculating the R^2^ and *p* value for Pearson’s correlation coefficient.

**Figure 3.**
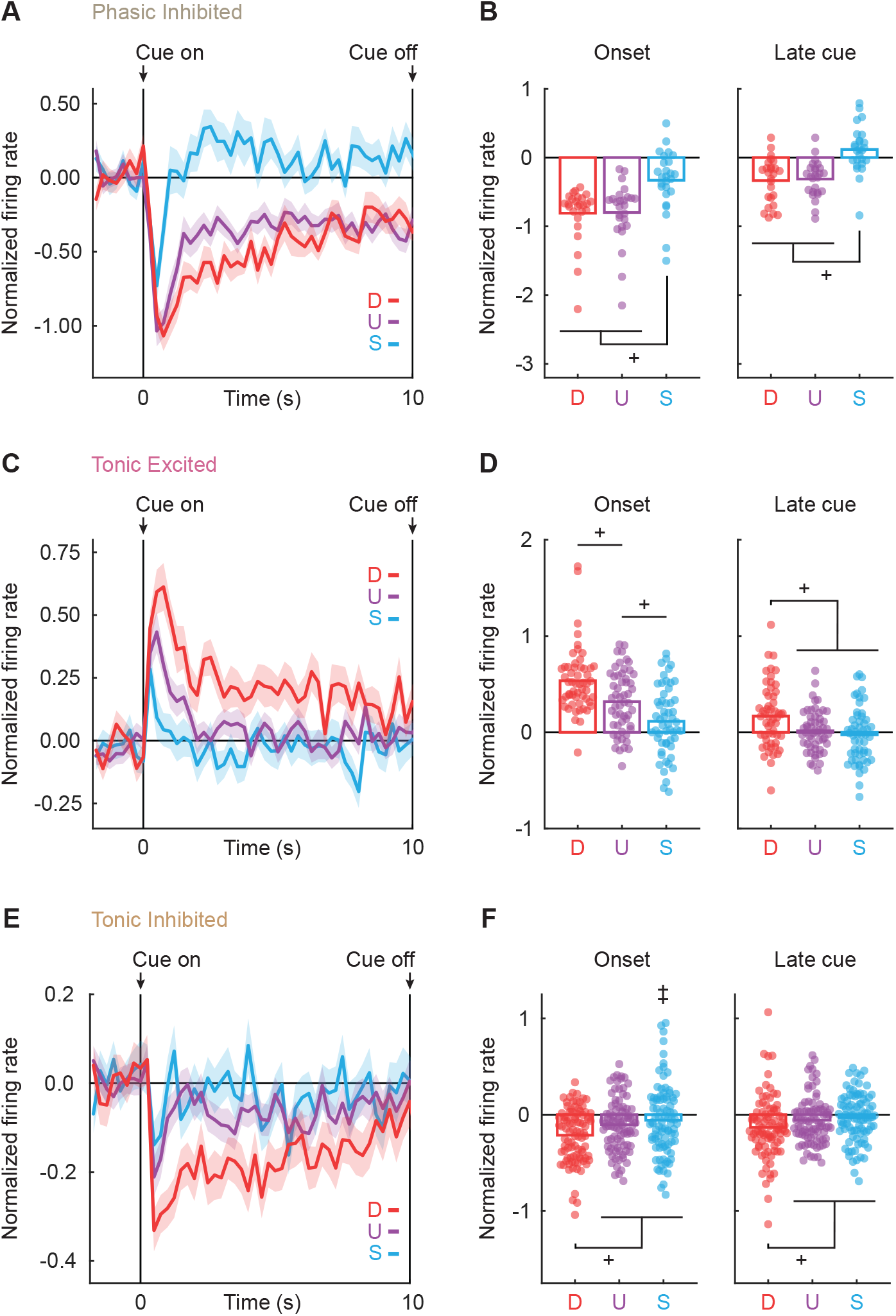
Differential firing in cue-responsive neurons. (**A**) Mean normalized firing rate to danger (D, red), uncertainty (U, purple), and safety (S, blue) is shown from 2 s prior to cue onset to cue offset for the Phasic Inhibited neurons (n = 26, grey). Cue onset and offset are indicated by vertical black lines. SEM is indicated by shading. (**B**) Mean (bar) and individual (data points), normalized firing rate for Phasic Inhibited neurons during the first 1-s cue interval (onset, left) and the last 5-s cue interval (late cue, right) are shown for each cue (D, danger; U, uncertainty; S, safety). Colors maintained from A. (**C**-**F**) Identical graphs made for Tonic Excited (n = 55, pink) and Tonic Inhibited neurons (n = 91, tan), as in A and B. ^+^95% bootstrap confidence interval for differential cue firing does not contain zero. ^‡^ Pitman-Morgan test, *p* < 0.05.

**Figure. 4.**
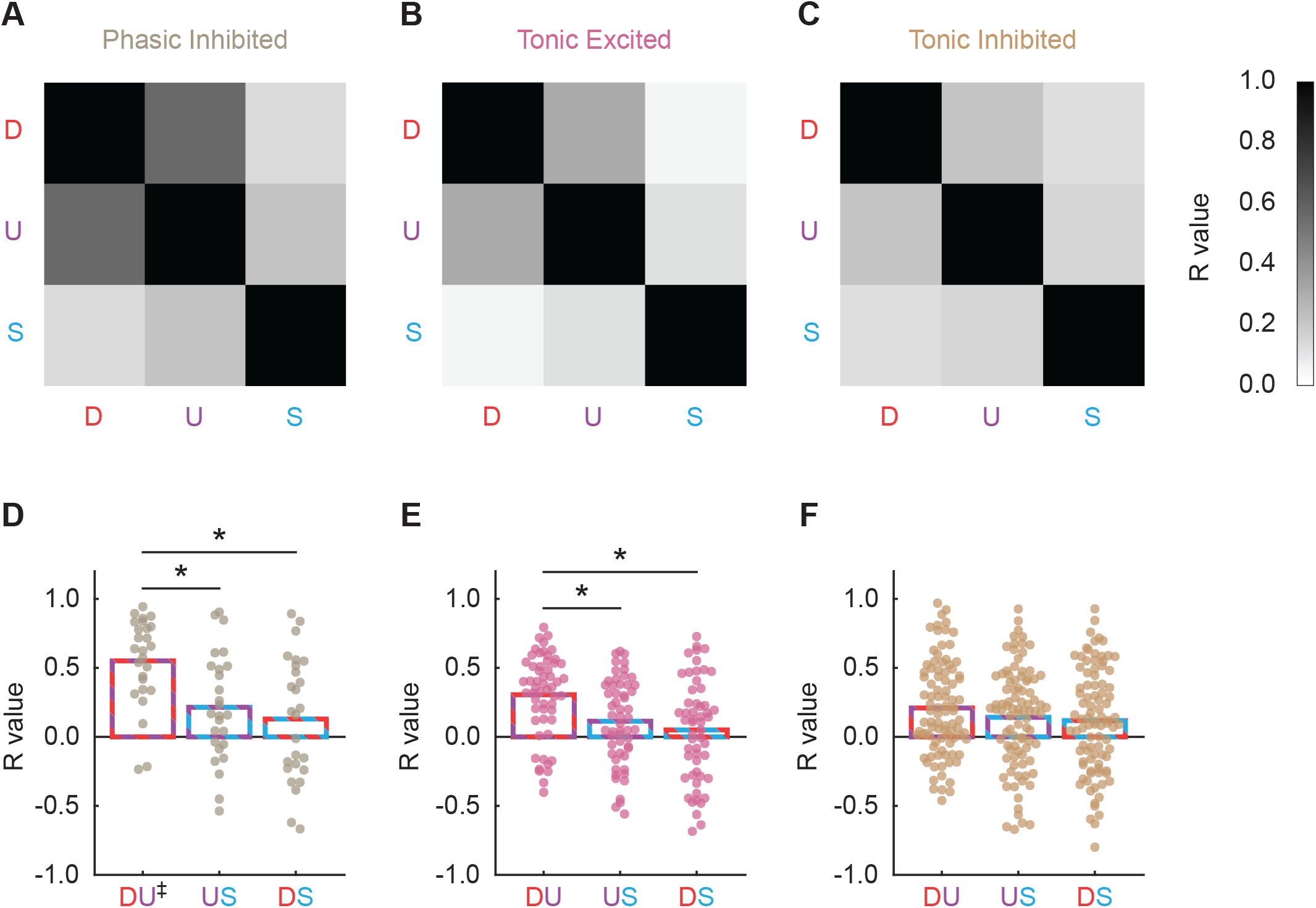
Firing similarity by cue-responsive neurons. (**A**-**C**) Firing similarity matrices between cue pairs (D, danger, red; U, uncertainty, purple; S, safety, blue) are depicted for (**A**) Phasic Inhibited (n = 26, grey), (**B**) Tonic Excited (n = 55, pink), and (**C**) Tonic Inhibited (n = 91, tan) neurons. Color scale for correlation coefficient (R) is shown to the right, with greatest firing similarities black (R = 1) and least firing similarities white (R = 0). (**D**-**F**) Mean (bar) and individual (data points) correlation coefficients (R) are shown for each cluster: (D-U) danger vs. uncertainty, (U-S) uncertainty vs. safety, and (D-S) danger vs. safety. *Bonferroni t-test, *p* < 0.05.

**Figure. 5.**
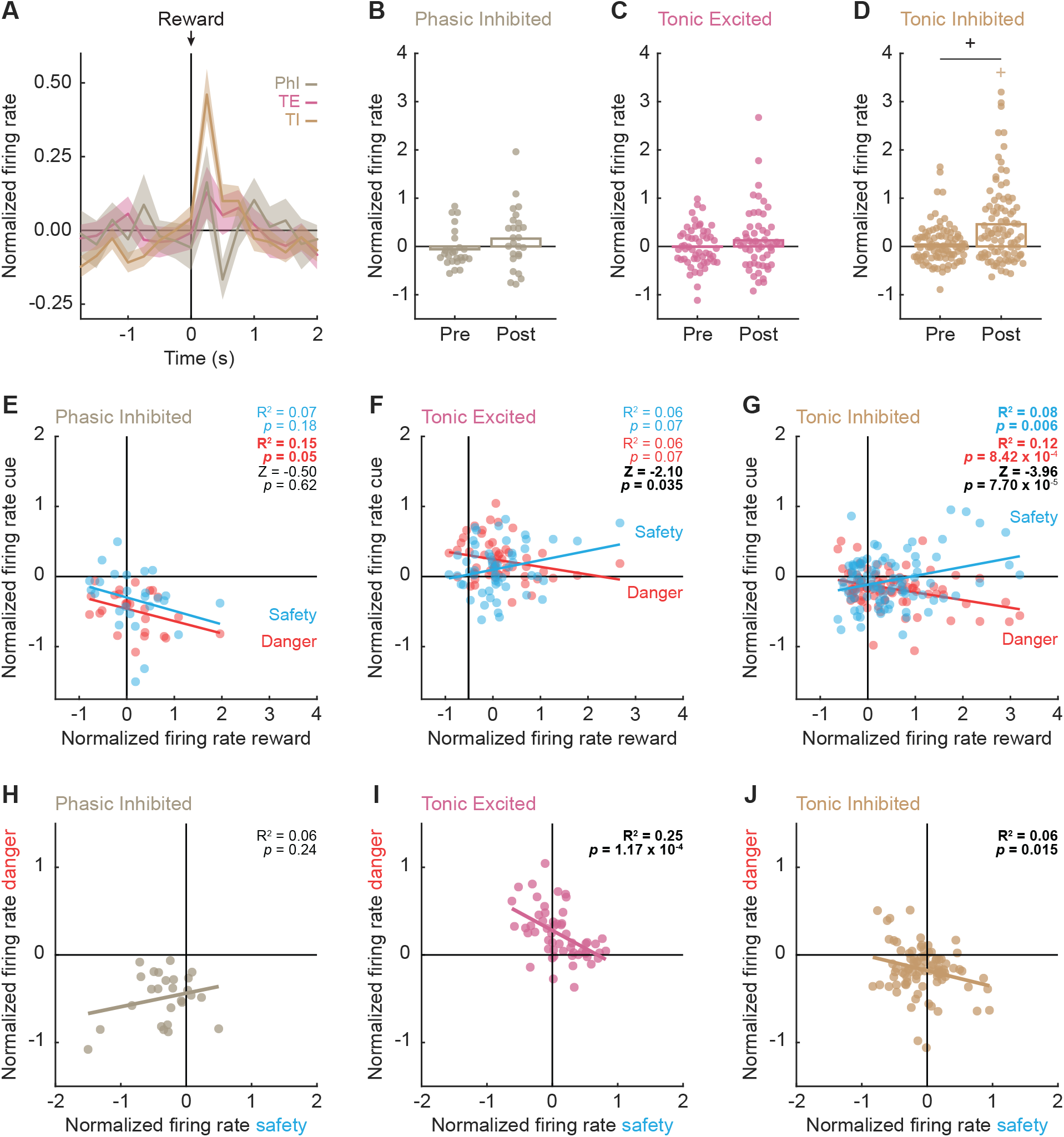
Tonic Inhibited neurons show opposing responses to danger and reward. (**A**) Mean ± SEM normalized firing rate to reward is shown 2 s prior to and 2 s after reward delivery (advancement of feeder) for the Phasic Inhibited (n = 26, grey), Tonic Excited (n = 55, pink), and Tonic Inhibited (n = 91, tan) neurons. Reward delivery is indicated by black arrow. SEM is indicated by shading. (**B**-**D**) Mean (bar) and individual (data points), normalized firing rate for (**B**) Phasic Inhibited, (**C**) Tonic Excited, and (**D**) Tonic Inhibited neurons are shown during 250-ms interval prior (pre) to and 250-ms interval after (post) reward delivery. ^+^95% bootstrap confidence interval for differential reward firing does not contain zero. ^+^95% bootstrap confidence interval for normalized firing rate does not contain zero (colored plus sign). (**E**-**G**) Mean normalized firing rate to cue (danger (10-s cue), red; Safety (the first 1-s cue), blue) vs. reward is plotted for (**E**) Phasic Inhibited, (**F**) Tonic Excited, and (**G**) Tonic Inhibited neurons. (**H**-**J**) Mean normalized firing rate to danger (10-s cue) vs. safety (the first 1-s cue) is plotted for (**H**) Phasic Inhibited, (**I**) Tonic Excited, and (**J**) Tonic Inhibited neurons. Trendline, the square of the Pearson correlation coefficient (R^2^) and associated *p* value are shown. Fisher r-to-z transformation (Z) are shown. Colors maintained from A.

### Pitman-Morgan test

The Pitman-Morgan test is used to compare within-cluster variance in cue firing (safety vs. danger; safety vs. uncertainty; Figure 3F). *P* values are reported for each test.

### Hartigan’s Dip Statistic

Hartigan’s Dip Statistic is used to test if the waveform peak-valley duration of each functional cluster had more than one mode in their distribution. *P* values are reported for each test, with *p* < 0.05 rejecting the null hypothesis that a distribution contained one mode, supporting the interpretation of a bimodal distribution.

### Firing similarity analyses

For each neuron, mean normalized firing rates were calculated for danger, uncertainty and safety (12, 1-s bins: 2 s prior to cue onset → cue offset). Firing similarity was quantified for each cue pair, for each neuron using Pearson’s correlation coefficient. The result was an R value for each neuron/cue pair: danger vs. uncertainty, uncertainty vs. safety and danger vs. safety. ANOVA for R value was performed with cluster and comparison as factors. Within and between-cluster post-hoc comparisons were performed with Bonferroni t-tests.

## Results

Female, Long Evans rats (n = 7) were moderately food deprived and trained to nose poke in a central port to receive food reward. Nose poking was reinforced throughout fear discrimination, but poke-reward contingencies were independent of cue-shock contingencies. During fear discrimination (Figure 1A, B), three auditory cues predicted unique foot shock probabilities: danger (*p* = 1.00), uncertainty (*p* = 0.25), and safety (*p* = 0.00). After eight discrimination sessions, rats were implanted with drivable, 16-wire microelectrode bundles dorsal to the NAcc. Following recovery, single-unit activity was recorded during fear discrimination. At the conclusion of recording, rats were perfused, brains sectioned, and electrode placements confirmed with Nissl staining (Figure 1C). Only placements within the NAcc, defined by a tear-shaped region surrounding the anterior commissure, were accepted (Figure 1D).

A total of 368 NAcc single units were recorded from 7 rats over 95 fear discrimination sessions. To identify cue-responsive single units in an unbiased manner, we compared mean baseline firing rate (Hz; 10 s prior to cue presentation) to mean firing rate during the first 1 s and last 5 s of cue presentation. A single unit was considered cue responsive if it showed a significant change (increase or decrease) in firing from baseline to danger, uncertainty, or safety during either the first 1-s or the last 5-s cue period (paired, two-tailed t-test, *p* < 0.05). This screen identified 193 cue-responsive single units (∼53% of all units recorded) from 82 sessions, with at least seven cue-responsive single units identified in each rat (Figure 1-1). All remaining analyses focus on cue-responsive NAcc single units (n = 193) and the fear discrimination sessions (n = 82) in which they were recorded.

Rats showed complete discrimination during the sessions in which cue-responsive single units were recorded (Figure 1E). Suppression ratios were high to danger, intermediate to uncertainty, and low to safety. ANOVA for mean individual suppression ratio revealed a main effect of cue (F_2,12_ = 57.41, *p* = 7.18 × 10^−7^, partial eta squared (η_p_^2^) = 0.91, observed power (op) = 1.00). Suppression ratios differed for each cue pair. The 95% bootstrap confidence interval for differential suppression ratio did not contain zero for danger vs. uncertainty (mean = 0.21, 95% CI [(lower bound) 0.16, (upper bound) 0.29]), uncertainty vs. safety (M = 0.47, 95% CI [0.35, 1.03]), and danger vs. safety (M = 0.69, 95% CI [0.50, 1.25]). Observing complete fear discrimination permits a meaningful examination of threat-related NAcc firing.

### NAcc neurons show heterogeneous cue responding

The firing pattern of cue-responsive NAcc single units varied considerably, as did the direction and magnitude of their response. Firing heterogeneity indicated that NAcc single units could be divided into discrete, functional populations. To identify these populations, we summarized firing in a 193 single unit x 6 epoch matrix. The six epochs were mean normalized firing rates taken from 2 periods (first 1 s and last 5 s) for each of the 3 cues (danger, uncertainty and safety). We applied k-means clustering to the matrix and found that five clusters grouped similar functional types (Figure 2-1).

To visualize cue firing patterns, we organized single units by cluster and plotted mean single-unit danger, uncertainty, safety, and reward firing (Figure 2A). Single units from two clusters showed strong phasic firing to danger and uncertainty, and lesser firing to safety: Phasic Excited neurons (n = 5; Figure 2A, Row 1), and Phasic Inhibited neurons (n = 26; Figure 2A, Row 2). Single units from two additional clusters showed modest tonic firing to danger, but lesser and similar firing to uncertainty and safety: Tonic Excited neurons (n = 55; Figure 2A, Row 3), and Tonic Inhibited neurons (n = 91; Figure 2A, Row 4). The final cluster (Non Selective, n = 16; Figure 2A, Row 5) showed firing increases that did not differentiate the cues. This was confirmed by ANOVA for normalized firing rate [factors: cue (danger, uncertainty, and safety) and interval (250-ms bins from 2 s prior to cue onset to cue offset)] which found neither a main effect of cue (F_2,30_ = 2.60, *p* = 0.091, η_p_^2^ = 0.15, op = 0.48) nor a cue x interval interaction (F_110,1650_ = 1.19, *p* = 0.098, η_p_^2^ = 0.07, op = 1.00).

Phasic Excited neurons (n = 5) were observed in only 2 of 7 subjects, with 4 single units coming from one subject. Thus, we are not confident these single units are representative of the NAcc. Phasic Inhibited neurons were obtained from 4 of 7 subjects, Tonic Excited neurons from 6 of 7 subjects and Tonic Inhibited neurons from all 7 subjects. Thus, the remaining analyses focused on NAcc-representative populations showing differential cue firing: Phasic Inhibited, Tonic Inhibited, and Tonic Excited neurons. Phasic Excited and Non-Selective (Figure 2-2) neuron analyses are provided as Extended Data.

In line with previous studies classifying single units based on firing characteristics, we sought to determine if our functionally defined clusters belonged to different neuron types (or a specific population of NAcc neurons). Our baseline firing rate was taken from a time period when rats were poking for and consuming food rewards – making it a poor indicator of firing at rest. We therefore focused on waveform peak-valley duration, which would be much less contaminated by behavior and has previously been shown to separate NAcc neuron types. Narrow waveforms are common in NAcc fast-spiking interneurons (FSIs), while wide waveforms are common in projecting, NAcc medium spiny neurons (MSNs) (Berke, 2008; Sosa et al., 2020; Gage, 2010; Gitts, 2011; Berke, 2004; Yarom, 2011; Vachez, 2021). Therefore, we calculated waveform peak-valley duration for each single unit. Consistent with recordings coming from a mix of FSIs and MSNs, we observed a bimodal distribution for waveform peak-valley duration across all clusters (Hartigan’s Dip Statistic, test could not provide specific *p* value, instead returned 0.00). Single units tended to fall at the histogram extremes: narrow or wide. We then applied Hartigan’s Dip Statistic to each cluster, determining whether it was primarily composed of one neuron type, or a mix of both.

Wide waveforms, putative MSNs, were more common among Phasic Inhibited neurons (Figure 2B). A bimodal distribution, indicating a mix of MSNs and FSIs, was only detected if an uncorrected *p* value was used (Hartigan’s Dip Statistic, *p* = 0.036). Tonic Excited neurons (Figure 2C) showed a unimodal distribution (Hartigan’s Dip Statistic *p* = 0.17), with most single units exhibiting narrow waveforms, typical of FSIs. Finally, Tonic Inhibited neurons (Figure 2D) showed a bimodal distribution (Hartigan’s Dip Statistic, test could not provide specific *p* value, instead returned 0.00), with almost half of the single units exhibiting narrow and the other half exhibiting wide waveforms (a mixture of putative MSNs and FSIs). Single-unit function is likely related to neuron type; an observation we return to in the discussion.

### Phasic Inhibited NAcc neurons are threat responsive

We have previously shown that NAcc activity during cue presentation is necessary to rapidly discriminate uncertainty and safety (Ray et al., 2020). We were curious whether phasic NAcc firing decreases rapidly discriminated threat cues (danger and uncertainty) from safety. To determine this, we performed ANOVA for normalized firing rate by Phasic Inhibited neurons (n = 26) [factors: cue (danger, uncertainty, and safety) and interval (250-ms bins, from 2 s prior to cue onset to cue offset)]. Confirming phasic and differential firing, ANOVA revealed a main effect of cue (F_2,50_ = 37.84, *p* = 9.83 × 10^−11^, η_p_^2^ = 0.60, op = 1.00), interval (F_47,1175_ = 11.51, *p* = 8.23 × 10^−68^, η_p_^2^ = 0.32, op = 1.00), and a significant cue x interval interaction (F_94,2350_ = 4.95, *p* = 7.61 × 10^−45^, η_p_^2^ = 0.17, op = 1.00). Population activity suggests equivalent firing decreases to danger and uncertainty that exceeded firing decreases to safety (Figure 3A). In support, Phasic Inhibited neurons showed greater firing decreases to threat cues (danger and uncertainty) compared to safety at onset (M = -0.47, 95% CI [-0.63, -0.32]; Figure 3B, left), and during late cue (M = -0.44, 95% CI [-0.60, -0.20]; Figure 3B, right). Phasic Inhibited neurons rapidly discriminated threat from safety.

### Tonic NAcc neurons are predominantly danger responsive

Our previous study also found that pre-training NAcc lesions disrupted fear discrimination across cue presentation (Ray et al., 2020). This was driven in part by reduced nose poke suppression to danger in NAcc-lesioned rats. Given that tonic neurons sustained firing over cue presentation (Figure 3C, E) we were curious whether these neurons showed differential cue firing that was more specific to danger. Confirming differential firing for Tonic Excited neurons (n = 55), ANOVA revealed a main effect of cue (F_2,106_ = 13.05, *p* = 9.00 × 10^−6^, η_p_^2^ = 0.20, op = 1.00), interval (F_47,2491_ = 7.12, *p* = 1.17 × 10^−41^, η_p_^2^ = 0.12, op = 1.00), and a significant cue x interval interaction (F_94,4982_ = 2.23, *p* = 1.87 × 10^−10^, η_p_^2^ = 0.04, op = 1.00). Firing was maximal to danger and fully discriminated the three cues at onset (danger vs. uncertainty: M = 0.22, 95% CI [0.11, 0.33]; uncertainty vs. safety (M = 0.21, 95% CI [0.07, 0.33]; Figure 3D, left). As cue presentation proceeded, firing increases were selective to danger whereas uncertainty and safety firing were minimal and equivalent. In support, Tonic Excited neurons showed differential firing to danger compared to the mean of uncertainty and safety during late cue presentation (M = 0.17, 95% CI [0.07, 0.27]; Figure 3D, right). Tonic Excited neuronal firing initially discriminated all cues before becoming specific to danger.

Tonic Inhibited neurons (n = 91) were the most abundant functional type, accounting for ∼47% of cue-responsive NAcc neurons. To reveal if these neurons also showed differential firing that was more specific to danger, we first performed ANOVA. Confirming differential firing, ANOVA revealed a significant main effect of cue (F_2,178_ = 8.90, *p* = 2.07 × 10^−4^, η_p_^2^ = 0.09, op = 0.97), interval (F_47,4183_ = 3.98, *p* = 4.49 × 10^−18^, η_p_ ^2^ = 0.04, op = 1.00), and a significant cue x interval interaction (F_94,8386_ = 1.45, *p* = 0.003, η_p_^2^ = 0.02, op = 1.00). Confirming more selective danger firing, Tonic Inhibited neurons showed danger firing decreases that exceeded uncertainty and safety decreases at onset (M = -0.13, 95% CI [-0.22, -0.03]; Figure 3F, left), as well as late cue (M = -0.09, 95% CI [-0.16, -0.02]; Figure 3F, right).

As a population, Tonic Inhibited neurons showed minimal safety firing. However, inspection of the individual units (Figure 3F, left) revealed considerable variation in safety onset firing, with some neurons showing large safety firing increases. Supporting this observation, there was greater variability in onset safety firing compared to either danger (Pitman-Morgan test, *p* = 0.0087) or uncertainty (Pitman-Morgan test, *p* = 0.0046). Tonic Inhibited neuronal firing distinguished danger from uncertainty and safety throughout cue presentation, though neurons varied in their safety onset firing.

### Phasic Inhibited neurons fire similarly to danger and uncertainty

Previous analyses reveal that Phasic Inhibited neurons fire more similarly to danger and uncertainty than Tonic Excited and Tonic Inhibited neurons. Perhaps Phasic Inhibited neurons also show more similar temporal firing patterns to danger and uncertainty over the duration of cue presentation. Inspired by representational similarity analyses (Kriegeskorte et al., 2008; Ritchey et al., 2013), we devised a firing similarity analysis. To do this, we divided mean normalized firing rate for each cue into 12, 1-s bins (2 s prior to cue onset through 10-s cue presentation). We correlated firing rate over this 12-s period for each neuron/cue pair using Pearson’s correlation coefficient. Firing similarity for each cluster was determined using the mean R value for each neuron/cue pair (Figure 4 A-C).

Phasic Inhibited neurons showed greatest firing similarity to danger and uncertainty (Figure 4D). Danger/uncertainty firing similarity eclipsed similarity for uncertainty/safety and danger/safety. Further, danger/uncertainty firing similarity by Phasic Inhibited neurons exceeded that for Tonic Excited (Figure 4E) and Tonic Inhibited neurons (Figure 4F). In support, ANOVA for R value [factors: cluster (Phasic Inhibited, Tonic Excited, and Tonic Inhibited) and comparison (danger/uncertainty, uncertainty/safety and danger/safety)] revealed significant main effects of cluster (F_2,169_ = 3.34, *p* = 0.038, η_p_^2^ = 0.04, op = 0.63) and comparison (F_2,338_ = 26.26, *p* = 2.51 × 10^−11^, η_p_^2^ = 0.13, op = 1.00), but critically a significant cluster x comparison interaction (F_4,338_ = 3.86, *p* = 0.004, η_p_^2^ = 0.04, op = 0.90). Paired t-tests (Bonferroni corrected for 9 tests; 0.05/9 = 0.0056) revealed Phasic Inhibited R values for danger/uncertainty to exceed those for uncertainty/safety (t_25_ = 3.31, *p* = 0.003) and danger/safety (t_25_ = 4.11, *p* = 3.74 × 10^−4^). Tonic Excited R values for danger/uncertainty exceed those for uncertainty/safety (t_54_ = 3.24, *p* = 0.002) and danger/safety (t_54_ = 4.11, *p* = 9.60 × 10^−5^); but no differences were observed for any cue pair in Tonic Inhibited neurons (all t < 2.1, all *p* > 0.0056). Across all clusters, greatest firing similarity was observed for danger and uncertainty firing by Phasic Inhibited neurons.

### Distinct NAcc signals for threat and valence

The goal of the current experiment was to examine NAcc threat-related firing. Of course, the NAcc is best known for its role in reward-related behavior. While our procedure was optimized to assess threat, the use of conditioned suppression permitted us to record activity around reward delivery. To examine if our cue-responsive populations showed reward-related responding, we aligned firing of our three main clusters (Phasic Inhibited, Tonic Excited, and Tonic Inhibited) to pellet feeder advance (Figure 5A). We performed ANOVA for normalized firing rate [factors: cluster (Phasic Inhibited, Tonic Excited, Tonic Inhibited) and interval (16, 250-ms bins: 2 s prior to and 2 s following pellet feeder advance)]. ANOVA revealed a main effect of interval (F_15,2430_ = 3.83, *p* = 8.49 × 10^−7^, η_p_^2^ = 0.02, op = 1.00), but more critically, a cluster x interval interaction (F_30,2430_ = 1.72, *p* = 0.009, η_p_^2^ = 0.02, op = 1.00). The interaction was the result of Tonic Inhibited neurons selectively increasing firing following pellet feeder advance. ANOVA restricted to Tonic Inhibited neurons found a main effect of interval (F_15,1290_ = 8.78, *p* = 1.24 × 10^−19^, η_p_^2^ = 0.09, op = 1.00), while separate ANOVA for Tonic Excited and Phasic Inhibited neurons found no main effects of interval (F < 1.1, *p* > 0.4). Population firing patterns were evident in single units (Figure 5B-D). In Phasic Inhibited and Tonic Excited neurons, pre and post reward firing differed neither from zero nor from each other (all 95% CIs contained zero; Figure 5 B, C). By contrast, Tonic Inhibited firing around zero prior to reward (M = 0.04, 95% CI [-0.07, 0.13]) gave way to firing increases post reward (M = 0.47, 95% CI [0.28, 0.62]), and post reward firing exceeded pre reward firing (M = 0.42, 95% CI [0.21, 0.58]; Figure 5D).

Tonic Inhibited neurons – which increased firing following reward, sustained firing decreases to danger and showed variable firing increases to safety onset – may generally signal valence. If this were the case, positive firing relationships would be observed for safety onset and reward onset (both of which have positive valence); and negative firing relationships would be observed for danger and reward (which have opposing valence). Phasic Inhibited neurons (Figure 5E) showed a non-significant, negative firing relationship between reward onset firing and safety onset firing (R^2^ = 0.07, *p* = 0.18), a significant, negative firing relationship between reward onset firing and danger firing (R^2^ = 0.15, *p* = 0.05), and these two correlations did not differ from one another (Z = -0.50, *p* = 0.62). Tonic Excited neurons (Figure 5F) showed a non-significant positive firing relationship between reward onset firing and safety onset firing (R^2^ = 0.06, *p* = 0.07), a non-significant, negative firing relationship between reward onset firing and danger firing (R^2^ = 0.06, *p* = 0.07), but these two correlations differed from one another (Z = -2.10, *p* = 0.035). Thus, neither Phasic Inhibited nor Tonic Excited neurons are strong candidates for general valence signaling.

Tonic Inhibited neurons (Figure 5G) showed a significant, positive relationship between reward onset firing and safety onset firing (R^2^ = 0.08, *p* = 0.006), a significant, negative relationship between reward onset firing and danger firing (R^2^ = 0.12, *p* = 8.42 × 10^−4^), and these two correlations significantly differed from one another (Z = -3.96, *p* = 7.70 × 10^−5^). Finally, if neurons signal valence, opposing firing changes should be observed to safety onset and danger. While Phasic Inhibited neurons (R^2^ = 0.06, *p* = 0.24; Figure 5H) showed zero firing relationship, Tonic Excited (R^2^ = 0.25, *p* = 1.17 × 10^−4^; Figure 5I) and Tonic Inhibited (R^2^ = 0.06, *p* = 0.015; Figure 5J) neurons showed significant, negative firing relationships. The results reveal selective threat signaling by Phasic Inhibited neurons, somewhat ambiguous signaling by Tonic Excited neurons and complete valence signaling by Tonic Inhibited neurons.

## Discussion

We recorded NAcc single-unit activity while female rats discriminated danger, uncertainty, and safety. Demonstrating direct threat responding, most NAcc neurons showed greatest firing changes to the shock-associated cues: danger and uncertainty. Revealing diverse roles for the NAcc in threat and valence, heterogeneity in cue firing patterns led us to detect at least three functional types. These types included neurons responding to danger and uncertainty through firing decreases versus increases; and neurons that specifically signaled threat versus neurons that signaled valence through opposing changes in firing to negative valence (danger and uncertainty) and positive valence (reward and safety onset).

Before discussing the organization of threat and valence responding within the NAcc and its relationship to a larger network, consideration of biological sex is necessary. In a previous study we manipulated NAcc activity in only male rats via neurotoxic lesion or optogenetic inhibition (Ray et al., 2020). We found that the NAcc is necessary for the acquisition of fear to danger, as well as the acquisition of discriminative fear to threat and safety cues. We further found that NAcc activity at the time of cue presentation is necessary for expression of rapid discrimination of uncertainty and safety. To reveal NAcc function across biological sex, we devised hypotheses from our male-only manipulation data and tested hypotheses by recording only in females. Our female-only results mostly confirmed our male-derived hypotheses. Tonic NAcc responses discriminated all cues but biased firing to danger. Phasic NAcc responses separated threat (uncertainty and danger) from safety. The concordance of our male manipulation and female recording findings may be less surprising, given that our laboratory observes only modest sex differences in our variant of fear discrimination (Walker et al., 2018, 2019). Collectively, our findings suggest that NAcc threat function may be conserved across female and male adult rats.

Consistent with prior studies (Berke, 2008; Gage et al., 2010; Lansink et al., 2010; Sosa et al., 2020; Vachez et al., 2021), we found that waveform duration was bimodally distributed in cue-responsive NAcc neurons. Narrow waveforms are consistent with FSIs, reflecting firing information confined within the NAcc. Wide waveforms are consistent with MSNs, the principal output neurons of the NAcc. Although one might expect tight coupling of FSI and MSN firing patterns *in vivo*, this has not been consistently observed (Lansink et al., 2010; Berke, 2011). Moreover, firing of individual FSIs is often idiosyncratic and unrelated to neighboring neurons. We found that putative FSIs were a mix of responses to danger through firing excitation and firing inhibition. Putative FSIs further varied in the duration of their danger response (phasic vs. tonic), the degree to which they responded following reward delivery, and their relationship between danger and reward firing. Even in threat situations, NAcc FSIs remain enigmatic.

By contrast, more consistent firing patterns were observed in putative MSNs. Wide-waveform NAcc neurons universally suppressed firing during danger cue presentation. Two distinct signals emerged, with phasic neurons showing equivalent firing inhibition to danger and uncertainty; but tonic neurons showing preferential firing inhibition to danger. Thus, phasic neurons may signal threat and tonic neurons may signal danger. Might these functional types correspond to unique genetic neuron types? NAcc MSNs are primarily composed of two genetic types, GABA neurons expressing dopamine D1 vs. D2 receptors. It is tempting to speculate that tonic inhibited neurons are D1-MSNs. Rewarding stimuli are proposed to activate D1-MSNs more greatly than D2-MSNs (Hikida et al., 2016). NAcc D1 projections to VP drive cocaine seeking (Pardo-Garcia et al., 2019). Rats will optogenetically self-stimulate NAcc D1-MSNs at greater rates than D2-MSNs (Cole et al., 2018). However, the NAcc does not respect the traditional D1-direct vs. D2-direct pathway like other subregions of the striatum. Both NAcc D1- and D2-MSNs project to the ventral pallidum (Kupchik et al., 2015). Depending on the stimulation pattern, D1- and D2-MSNs can both drive approach and avoidance behaviors (Soares-Cunha et al., 2020). Even more, a small population of NAcc neurons appear to express both D1 and D2 receptors (Le Moine and Bloch, 1996; Lee et al., 2006). So, while tempting to speculate the genetic identity of our tonic inhibited neurons, any conclusion reached would be premature.

Getting threat information to the NAcc seems obvious, as it receives direct input from the basolateral amygdala. Less obvious is how NAcc threat information is output to a larger network, as there are no reciprocal projections from the NAcc to any amygdala subregion. Recent work from our laboratory suggest the ventral pallidum, which receives direct NAcc input, is a compelling candidate (Moaddab et al., 2021). Recording ventral pallidum single-unit activity in the same fear discrimination procedure, we revealed a large functional population signaling relative threat. That is, many ventral pallidum neurons linearly decrease cue firing according to foot shock probability. The ventral pallidum is anatomically linked to the amygdala, receiving inputs from the central amygdala, and projecting directly to the basolateral amygdala. NAcc threat signals may train up relative threat signaling in the ventral pallidum during acquisition of fear discrimination. An enduring role for the NAcc may be to rapidly classify safe vs. threat stimuli, consistent with rapid NAcc processing of reward (Richard et al., 2016; Ottenheimer et al., 2018).

Of course, the ventral pallidum is unlikely the sole recipient of NAcc threat information. The NAcc projects directly to the ventrolateral periaqueductal gray, which has long been implicated in fear output, but we are finding to more flexibility signal threat probability (Usuda et al., 1998; Wright and McDannald, 2019; Wright et al., 2019). The NAcc also projects directly to the retrorubral field, a midbrain region housing A8 dopamine neurons (Deutch et al., 1988) that contains a diverse signals for threat and aversive outcome (Usuda et al., 1998; Moaddab and McDannald, 2021). These direct projections remain unstudied, and we are therefore only beginning to understand how NAcc threat responding may shape function and responding of a larger brain network.

We set to reveal NAcc threat responding. At the same time, we were able to record NAcc activity around reward delivery – a more familiar setting for investigations of NAcc function. We found many neurons defined with respect to threat showed reward responding. Specifically, a population of neurons that increased firing to reward *inhibited* firing to danger. This kind of bi-directional signaling is reminiscent of valence signals observed in the basolateral amygdala. Only in the basolateral amygdala, positive and negative valence signals are most commonly reported in distinct populations (Schoenbaum et al., 1999; Paton et al., 2006; Beyeler et al., 2016; Kim et al., 2016). This in includes basolateral amygdala neurons that show similar firing to reward and safety (Sangha et al., 2013). The NAcc may be a site in which parallel inputs for positive and negative valence from the basolateral amygdala converge. Like the NAcc, the ventral pallidum contains a large, functional population that shows bi-directional firing to threat (firing inhibition) and reward (firing increase) (Kaplan et al., 2020; Stephenson-Jones et al., 2020; Moaddab et al., 2021). Bi-directional signaling of valence may be an organizational feature of ventral striatum and ventral pallidum.

Our findings of robust, diverse signaling of threat within the NAcc is supported by clinical observations of NAcc volume and activity relationships with anxiety. Larger NAcc volume is associated with higher social anxiety (Günther et al., 2018), as well as higher trait anxiety (Kühn et al., 2011). NAcc volume has also been shown to predict anxiety treatment success (Burkhouse et al., 2020). Viewing traumatic images elicits a NAcc BOLD response (Liberzon et al., 1999). These results suggest NAcc may be linked to anxiety dysfunction as well as successful treatment. Although correlative, this literature suggests the NAcc normally plays a normal role in threat behavior and may be a critical site of disruption in stress and anxiety disorders. Our results provide a framework for threat and valence processing within the NAcc, working to more fully map neural circuits for normal and disrupted threat function.

## Data and software availability

Full electrophysiology data set will be uploaded to http://crcns.org/ upon acceptance for publication.

## Additional resources

Med Associates programs used for behavior and Matlab programs used for behavioral analyses are made freely available at our lab website: http://mcdannaldlab.org/resources

## Acknowledgements

We thank Bret Judson and the Boston College Imaging Core for infrastructure/support, and Dr. Maureen Ritchey for constructive discussions related to the project.

## References

Ambroggi F, Ghazizadeh A, Nicola SM, Fields HL (2011) Roles of nucleus accumbens core and shell in incentive-cue responding and behavioral inhibition. J Neurosci 31:6820–6830.

Badrinarayan A, Wescott SA, Vander Weele CM, Saunders BT, Couturier BE, Maren S, Aragona BJ (2012) Aversive Stimuli Differentially Modulate Real-Time Dopamine Transmission Dynamics within the Nucleus Accumbens Core and Shell. Journal of Neuroscience 32:15779–15790.

Baldo BA, Kelley AE (2007) Discrete neurochemical coding of distinguishable motivational processes: insights from nucleus accumbens control of feeding. Psychopharmacology 191:439–459.

Basar K, Sesia T, Groenewegen H, Steinbusch HWM, Visser-Vandewalle V, Temel Y (2010) Nucleus accumbens and impulsivity. Progress in Neurobiology 92:533–557.

Beck CH, Fibiger HC (1995) Conditioned fear-induced changes in behavior and in the expression of the immediate early gene c-fos: with and without diazepam pretreatment. Journal of Neuroscience 15:709–720.

Berke JD (2008) Uncoordinated Firing Rate Changes of Striatal Fast-Spiking Interneurons during Behavioral Task Performance. J Neurosci 28:10075–10080.

Berke JD (2011) Functional Properties of Striatal Fast-Spiking Interneurons. Front Syst Neurosci 5 Available at: https://www.frontiersin.org/articles/10.3389/fnsys.2011.00045/full [Accessed May 13, 2021].

Berridge KC (2019) Affective valence in the brain: modules or modes? Nat Rev Neurosci 20:225–234.

Beyeler A, Namburi P, Glober GF, Simonnet C, Calhoon GG, Conyers GF, Luck R, Wildes CP, Tye KM (2016) Divergent Routing of Positive and Negative Information from the Amygdala during Memory Retrieval. Neuron 90:348–361.

Blaiss CA, Janak PH (2009) The nucleus accumbens core and shell are critical for the expression, but not the consolidation, of Pavlovian conditioned approach. Behav Brain Res 200:22–32.

Bolles RC (1970) Species-Specific Defense Reactions and Avoidance Learning. Psychol Rev 77:32–48.

Bolles RC, Collier AC (1976) The effect of predictive cues on freezing in rats. Animal Learning & Behavior 4:6–8.

Bouchet CA, Miner MA, Loetz EC, Rosberg AJ, Hake HS, Farmer CE, Ostrovskyy M, Gray N, Greenwood BN (2018) Activation of Nigrostriatal Dopamine Neurons during Fear Extinction Prevents the Renewal of Fear. Neuropsychopharmacology 43:665–672.

Bouton ME, Bolles RC (1980) Conditioned fear assessed by freezing and by the suppression of three different baselines. Animal Learning & Behavior 8:429–434.

Brog JS, Salyapongse A, Deutch AY, Zahm DS (1993) The patterns of afferent innervation of the core and shell in the ?Accumbens? part of the rat ventral striatum: Immunohistochemical detection of retrogradely transported fluoro-gold. J Comp Neurol 338:255–278.

Budygin EA, Park J, Bass CE, Grinevich VP, Bonin KD, Wightman RM (2012) Aversive stimulus differentially triggers subsecond dopamine release in reward regions. Neuroscience 201:331–337.

Burkhouse KL, Jagan Jimmy, Defelice N, Klumpp H, Ajilore O, Hosseini B, Fitzgerald KD, Monk CS, Phan KL (2020) Nucleus accumbens volume as a predictor of anxiety symptom improvement following CBT and SSRI treatment in two independent samples. Neuropsychopharmacology 45:561–569.

Cai LX, Pizano K, Gundersen GW, Hayes CL, Fleming WT, Holt S, Cox JM, Witten IB (2020) Distinct signals in medial and lateral VTA dopamine neurons modulate fear extinction at different times. eLife 9:e54936.

Campeau S, Davis M (1995) Involvement of the central nucleus and basolateral complex of the amygdala in fear conditioning measured with fear-potentiated startle in rats trained concurrently with auditory and visual conditioned stimuli. J Neurosci 15:2301–2311.

Carlezon WA, Thomas MJ (2009) Biological substrates of reward and aversion: A nucleus accumbens activity hypothesis. Neuropharmacology 56:122–132.

Cerri DH, Saddoris MP, Carelli RM (2014) Nucleus accumbens core neurons encode value-independent associations necessary for sensory preconditioning. Behavioral Neuroscience 128:567–578.

Chaudhri N, Sahuque LL, Schairer WW, Janak PH (2010) Separable Roles of the Nucleus Accumbens Core and Shell in Context- and Cue-Induced Alcohol-Seeking. Neuropsychopharmacol 35:783–791.

Christie MJ, Summers RJ, Stephenson JA, Cook CJ, Beart PM (1987) Excitatory amino acid projections to the nucleus accumbens septi in the rat: A retrograde transport study utilizingd[3H]aspartate and [3H]GABA. Neuroscience 22:425–439.

Cole SL, Robinson MJF, Berridge KC (2018) Optogenetic self-stimulation in the nucleus accumbens: D1 reward versus D2 ambivalence. PLoS One 13:e0207694.

Corbit LH, Balleine BW (2011) The general and outcome-specific forms of pavlovian-instrumental transfer are differentially mediated by the nucleus accumbens core and shell. J Neurosci 31:11786–11794.

Crameri F (2018) Scientific colour maps (Version 4.0.0).

Deutch AY, Goldstein M, Baldino F Jr, Roth RH (1988) Telencephalic projections of the A8 dopamine cell group. Ann N Y Acad Sci 537:27–50.

Di Ciano P, Robbins TW, Everitt BJ (2008) Differential effects of nucleus accumbens core, shell, or dorsal striatal inactivations on the persistence, reacquisition, or reinstatement of responding for a drug-paired conditioned reinforcer. Neuropsychopharmacology 33:1413–1425.

Floresco SB, McLaughlin RJ, Haluk DM (2008) Opposing roles for the nucleus accumbens core and shell in cue-induced reinstatement of food-seeking behavior. Neuroscience 154:877–884.

Fraser KM, Janak PH (2017) Long-lasting contribution of dopamine in the nucleus accumbens core, but not dorsal lateral striatum, to sign-tracking. Eur J Neurosci 46:2047–2055.

Gage GJ, Stoetzner CR, Wiltschko AB, Berke JD (2010) Selective Activation of Striatal Fast-Spiking Interneurons during Choice Execution. Neuron 67:466–479.

Gittis AH, Leventhal DK, Fensterheim BA, Pettibone JR, Berke JD, Kreitzer AC (2011) Selective Inhibition of Striatal Fast-Spiking Interneurons Causes Dyskinesias. Journal of Neuroscience 31:15727–15731.

Grill HJ, Norgren R (1978) The taste reactivity test. I. Mimetic responses to gustatory stimuli in neurologically normal rats. Brain Research 143:263–279.

Groessl F, Munsch T, Meis S, Griessner J, Kaczanowska J, Pliota P, Kargl D, Badurek S, Kraitsy K, Rassoulpour A, Zuber J, Lessmann V, Haubensak W (2018) Dorsal tegmental dopamine neurons gate associative learning of fear. Nat Neurosci 21:952–962.

Günther V, Ihme K, Kersting A, Hoffmann K-T, Lobsien D, Suslow T (2018) Volumetric Associations Between Amygdala, Nucleus Accumbens, and Socially Anxious Tendencies in Healthy Women. Neuroscience 374:25–32.

Hikida T, Morita M, Macpherson T (2016) Neural mechanisms of the nucleus accumbens circuit in reward and aversive learning. Neuroscience Research 108:1–5.

Iordanova MD, Westbrook RF, Killcross AS (2006) Dopamine activity in the nucleus accumbens modulates blocking in fear conditioning. Eur J Neurosci 24:3265–3270.

Kaplan A, Mizrahi-Kliger AD, Israel Z, Adler A, Bergman H (2020) Dissociable roles of ventral pallidum neurons in the basal ganglia reinforcement learning network. Nature Neuroscience 23:556–564.

Kerfoot EC, Agarwal I, Lee HJ, Holland PC (2007) Control of appetitive and aversive taste-reactivity responses by an auditory conditioned stimulus in a devaluation task: a FOS and behavioral analysis. Learn Mem 14:581–589.

Killcross S, Robbins TW, Everitt BJ (1997) Different types of fear-conditioned behaviour mediated by separate nuclei within amygdala. Nature 388:377–380.

Kim J, Pignatelli M, Xu S, Itohara S, Tonegawa S (2016) Antagonistic negative and positive neurons of the basolateral amygdala. Nat Neurosci 19:1636–1646.

Klawonn AM, Malenka RC (2018) Nucleus Accumbens Modulation in Reward and Aversion. Cold Spring Harb Symp Quant Biol 83:119–129.

Koo JW, Han JS, Kim JJ (2004) Selective neurotoxic lesions of basolateral and central nuclei of the amygdala produce differential effects on fear conditioning. Journal of Neuroscience 24:7654–7662.

Krause M, German PW, Taha SA, Fields HL (2010) A Pause in Nucleus Accumbens Neuron Firing Is Required to Initiate and Maintain Feeding. Journal of Neuroscience 30:4746– 4756.

Kriegeskorte N, Mur M, Bandettini PA (2008) Representational similarity analysis -connecting the branches of systems neuroscience. Front Syst Neurosci 2 Available at: https://www.frontiersin.org/articles/10.3389/neuro.06.004.2008/full?utm_source=FWEB&utm_medium=NBLOG&utm_campaign=ECO_10YA_top-research [Accessed May 11, 2021].

Kühn S, Schubert F, Gallinat J (2011) Structural correlates of trait anxiety: Reduced thickness in medial orbitofrontal cortex accompanied by volume increase in nucleus accumbens. Journal of Affective Disorders 134:315–319.

Kupchik YM, Brown RM, Heinsbroek JA, Lobo MK, Schwartz DJ, Kalivas PW (2015) Coding the direct/indirect pathways by D1 and D2 receptors is not valid for accumbens projections. Nat Neurosci 18:1230-+.

Lansink CS, Goltstein PM, Lankelma JV, Pennartz CMA (2010) Fast-spiking interneurons of the rat ventral striatum: temporal coordination of activity with principal cells and responsiveness to reward. European Journal of Neuroscience 32:494–508.

Le Moine C, Bloch B (1996) Expression of the D3 dopamine receptor in peptidergic neurons of the nucleus accumbens: comparison with the D1 and D2 dopamine receptors. Neuroscience 73:131–143.

LeDoux JE, Cicchetti P, Xagoraris A, Romanski LM (1990) The lateral amygdaloid nucleus: sensory interface of the amygdala in fear conditioning. Journal of Neuroscience 10:1062–1069.

Lee K-W, Kim Y, Kim AM, Helmin K, Nairn AC, Greengard P (2006) Cocaine-induced dendritic spine formation in D1 and D2 dopamine receptor-containing medium spiny neurons in nucleus accumbens. Proc Natl Acad Sci U S A 103:3399–3404.

Levita L, Dalley JW, Robbins TW (2002) Disruption of Pavlovian contextual conditioning by excitotoxic lesions of the nucleus accumbens core. Behavioral Neuroscience 116:539– 552.

Li Z, Chen Z, Fan G, Li A, Yuan J, Xu T (2018) Cell-Type-Specific Afferent Innervation of the Nucleus Accumbens Core and Shell. Front Neuroanat 12:84.

Liberzon I, Taylor SF, Amdur R, Jung TD, Chamberlain KR, Minoshima S, Koeppe RA, Fig LM (1999) Brain activation in PTSD in response to trauma-related stimuli. Biological Psychiatry 45:817–826.

Mannella F, Gurney K, Baldassarre G (2013) The nucleus accumbens as a nexus between values and goals in goal-directed behavior: a review and a new hypothesis. Front Behav Neurosci Available at: http://journal.frontiersin.org/article/10.3389/fnbeh.2013.00135/abstract [Accessed April 27, 2021].

Martinez RCR, Oliveira AR, Macedo CE, Molina VA, Brandão ML (2008) Involvement of dopaminergic mechanisms in the nucleus accumbens core and shell subregions in the expression of fear conditioning. Neuroscience Letters 446:112–116.

McGinty VB, Lardeux S, Taha SA, Kim JJ, Nicola SM (2013) Invigoration of reward seeking by cue and proximity encoding in the nucleus accumbens. Neuron 78:910–922.

Moaddab M, Hyland BI, Brown CH (2015) Oxytocin excites nucleus accumbens shell neurons in vivo. Molecular and Cellular Neuroscience 68:323–330.

Moaddab M, McDannald MA (2021) Retrorubral field is a hub for diverse threat and aversive outcome signals. Current Biology:S0960982221002992.

Moaddab M, Ray MH, McDannald MA (2021) Ventral pallidum neurons dynamically signal relative threat. Communications Biology 4:1–14.

Ottenheimer D, Richard JM, Janak PH (2018) Ventral pallidum encodes relative reward value earlier and more robustly than nucleus accumbens. Nat Commun 9 Available at: ://WOS:000447697100008.

Pardo-Garcia TR, Garcia-Keller C, Penaloza T, Richie CT, Pickel J, Hope BT, Harvey BK, Kalivas PW, Heinsbroek JA (2019) Ventral Pallidum Is the Primary Target for Accumbens D1 Projections Driving Cocaine Seeking. J Neurosci 39:2041–2051.

Parkinson JA, Olmstead MC, Burns LH, Robbins TW, Everitt BJ (1999a) Dissociation in effects of lesions of the nucleus accumbens core and shell on appetitive pavlovian approach behavior and the potentiation of conditioned reinforcement and locomotor activity by D-amphetamine. Journal of Neuroscience 19:2401–2411.

Parkinson JA, Robbins TW, Everitt BJ (1999b) Selective excitotoxic lesions of the nucleus accumbens core and shell differentially affect aversive Pavlovian conditioning to discrete and contextual cues. Psychobiology 27:256–266.

Parkinson JA, Willoughby PJ, Robbins TW, Everitt BJ (2000) Disconnection of the anterior cingulate cortex and nucleus accumbens core impairs Pavlovian approach behavior: further evidence for limbic cortical-ventral striatopallidal systems. Behav Neurosci 114:42–63.

Paton JJ, Belova MA, Morrison SE, Salzman CD (2006) The primate amygdala represents the positive and negative value of visual stimuli during learning. Nature 439:865–870.

Pauli WM, Larsen T, Collette S, Tyszka JM, Seymour B, O’Doherty JP (2015) Distinct Contributions of Ventromedial and Dorsolateral Subregions of the Human Substantia Nigra to Appetitive and Aversive Learning. Journal of Neuroscience 35:14220–14233.

Paxinos G, Watson C (2007) The rat brain in stereotaxic coordinates, 6th ed. Amsterdam ; Boston ; Academic Press/Elsevier. Available at: Publisher description http://www.loc.gov/catdir/enhancements/fy0745/2006937142-d.html.

Piantadosi PT, Yeates DCM, Floresco SB (2020) Prefrontal cortical and nucleus accumbens contributions to discriminative conditioned suppression of reward-seeking. Learn Mem 27:429–440.

Ray MH, Russ AN, Walker RA, McDannald MA (2020) The Nucleus Accumbens Core is Necessary to Scale Fear to Degree of Threat. J Neurosci 40:4750–4760.

Reynolds SM, Berridge KC (2002) Positive and Negative Motivation in Nucleus Accumbens Shell: Bivalent Rostrocaudal Gradients for GABA-Elicited Eating, Taste “Liking”/”Disliking” Reactions, Place Preference/Avoidance, and Fear. J Neurosci 22:7308–7320.

Richard JM, Ambroggi F, Janak PH, Fields HL (2016) Ventral Pallidum Neurons Encode Incentive Value and Promote Cue-Elicited Instrumental Actions. Neuron 90:1165–1173.

Ritchey M, Wing EA, LaBar KS, Cabeza R (2013) Neural Similarity Between Encoding and Retrieval is Related to Memory Via Hippocampal Interactions. Cerebral Cortex 23:2818– 2828.

Roesch MR, Singh T, Brown PL, Mullins SE, Schoenbaum G (2009) Ventral striatal neurons encode the value of the chosen action in rats deciding between differently delayed or sized rewards. J Neurosci 29:13365–13376.

Roitman MF, Wheeler RA, Carelli RM (2005) Nucleus accumbens neurons are innately tuned for rewarding and aversive taste stimuli, encode their predictors, and are linked to motor output. Neuron 45:587–597.

Saeb-Parsy K, Dyball REJ (2003) Defined Cell Groups in the Rat Suprachiasmatic Nucleus Have Different Day/Night Rhythms of Single-Unit Activity In Vivo. J Biol Rhythms 18:26–42.

Sangha S, Chadick JZ, Janak PH (2013) Safety encoding in the basal amygdala. J Neurosci 33:3744–3751.

Schoenbaum G, Chiba AA, Gallagher M (1999) Neural encoding in orbitofrontal cortex and basolateral amygdala during olfactory discrimination learning. Journal of Neuroscience 19:1876–1884.

Schoenbaum G, Setlow B (2003) Lesions of nucleus accumbens disrupt learning about aversive outcomes. J Neurosci 23:9833–9841.

Schwienbacher I, Fendt M, Richardson R, Schnitzler HU (2004) Temporary inactivation of the nucleus accumbens disrupts acquisition and expression of fear-potentiated startle in rats. Brain Res 1027:87–93.

Setlow B, Schoenbaum G, Gallagher M (2003) Neural encoding in ventral striatum during olfactory discrimination learning. Neuron 38:625–636.

Sicre M, Meffre J, Louber D, Ambroggi F (2020) The Nucleus Accumbens Core Is Necessary for Responding to Incentive But Not Instructive Stimuli. J Neurosci 40:1332–1343.

Soares-Cunha C, de Vasconcelos NAP, Coimbra B, Domingues AV, Silva JM, Loureiro-Campos E, Gaspar R, Sotiropoulos I, Sousa N, Rodrigues AJ (2020) Nucleus accumbens medium spiny neurons subtypes signal both reward and aversion. Molecular Psychiatry 25:3241–3255.

Sosa M, Joo HR, Frank LM (2020) Dorsal and Ventral Hippocampal Sharp-Wave Ripples Activate Distinct Nucleus Accumbens Networks. Neuron 105:725-741.e8.

Stephenson-Jones M, Bravo-Rivera C, Ahrens S, Furlan A, Xiao X, Fernandes-Henriques C, Li B (2020) Opposing Contributions of GABAergic and Glutamatergic Ventral Pallidal Neurons to Motivational Behaviors. Neuron 105:921-933.e5.

Sugam JA, Saddoris MP, Carelli RM (2014) Nucleus accumbens neurons track behavioral preferences and reward outcomes during risky decision making. Biol Psychiatry 75:807– 816.

Taha SA (2005) Encoding of Palatability and Appetitive Behaviors by Distinct Neuronal Populations in the Nucleus Accumbens. Journal of Neuroscience 25:1193–1202.

Thomas KL, Hall J, Everitt BJ (2002) Cellular imaging with zif268 expression in the rat nucleus accumbens and frontal cortex further dissociates the neural pathways activated following the retrieval of contextual and cued fear memory. European Journal of Neuroscience 16:1789–1796.

Usuda I, Tanaka K, Chiba T (1998) Efferent projections of the nucleus accumbens in the rat with special reference to subdivision of the nucleus: biotinylated dextran amine study. Brain Research 797:73–93.

Vachez YM, Tooley JR, Abiraman K, Matikainen-Ankney B, Casey E, Earnest T, Ramos LM, Silberberg H, Godynyuk E, Uddin O, Marconi L, Le Pichon CE, Creed MC (2021) Ventral arkypallidal neurons inhibit accumbal firing to promote reward consumption. Nat Neurosci 24:379–390.

Vetere G, Kenney JW, Tran LM, Xia F, Steadman PE, Parkinson J, Josselyn SA, Frankland PW (2017) Chemogenetic Interrogation of a Brain-wide Fear Memory Network in Mice. Neuron 94:363–374 e4.

Voorn P, Gerfen CR, Groenewegen HJ (1989) Compartmental organization of the ventral striatum of the rat: Immunohistochemical distribution of enkephalin, substance P, dopamine, and calcium-binding protein. J Comp Neurol 289:189–201.

Walker RA, Andreansky C, Ray MH, McDannald MA (2018) Early adolescent adversity inflates threat estimation in females and promotes alcohol use initiation in both sexes. Behav Neurosci 132:171–182.

Walker RA, Wright KM, Jhou TC, McDannald MA (2019) The ventrolateral periaqueductal gray updates fear via positive prediction error. Eur J Neurosci Available at: https://www.ncbi.nlm.nih.gov/pubmed/31376295.

Wendler E, Gaspar JCC, Ferreira TL, Barbiero JK, Andreatini R, Vital Mabf, Blaha Cd, Winn P, Da Cunha C (2014) The roles of the nucleus accumbens core, dorsomedial striatum, and dorsolateral striatum in learning: Performance and extinction of Pavlovian fear-conditioned responses and instrumental avoidance responses. Neurobiology of Learning and Memory 109:27–36.

Wenzel JM, Rauscher NA, Cheer JF, Oleson EB (2015) A Role for Phasic Dopamine Release within the Nucleus Accumbens in Encoding Aversion: A Review of the Neurochemical Literature. ACS Chem Neurosci 6:16–26.

Wright KM, Jhou TC, Pimpinelli D, McDannald MA (2019) Cue-inhibited ventrolateral periaqueductal gray neurons signal fear output and threat probability in male rats. Elife 8 Available at: https://www.ncbi.nlm.nih.gov/pubmed/31566567.

Wright KM, McDannald MA (2019) Ventrolateral periaqueductal gray neurons prioritize threat probability over fear output. Elife 8 Available at: https://www.ncbi.nlm.nih.gov/pubmed/30843787.

Zahm DS, Brog JS (1992) On the significance of subterritories in the “accumbens” part of the rat ventral striatum. Neuroscience 50:751–767.

Zhang Y, Zhu Y, Cao S-X, Sun P, Yang J-M, Xia Y-F, Xie S-Z, Yu X-D, Fu J-Y, Shen C-J, He H-Y, Pan H-Q, Chen X-J, Wang H, Li X-M (2020) MeCP2 in cholinergic interneurons of nucleus accumbens regulates fear learning. eLife 9:e55342.

