## Supplemental Figures for "Distinct threat and valence signals in rat nucleus accumbens core"

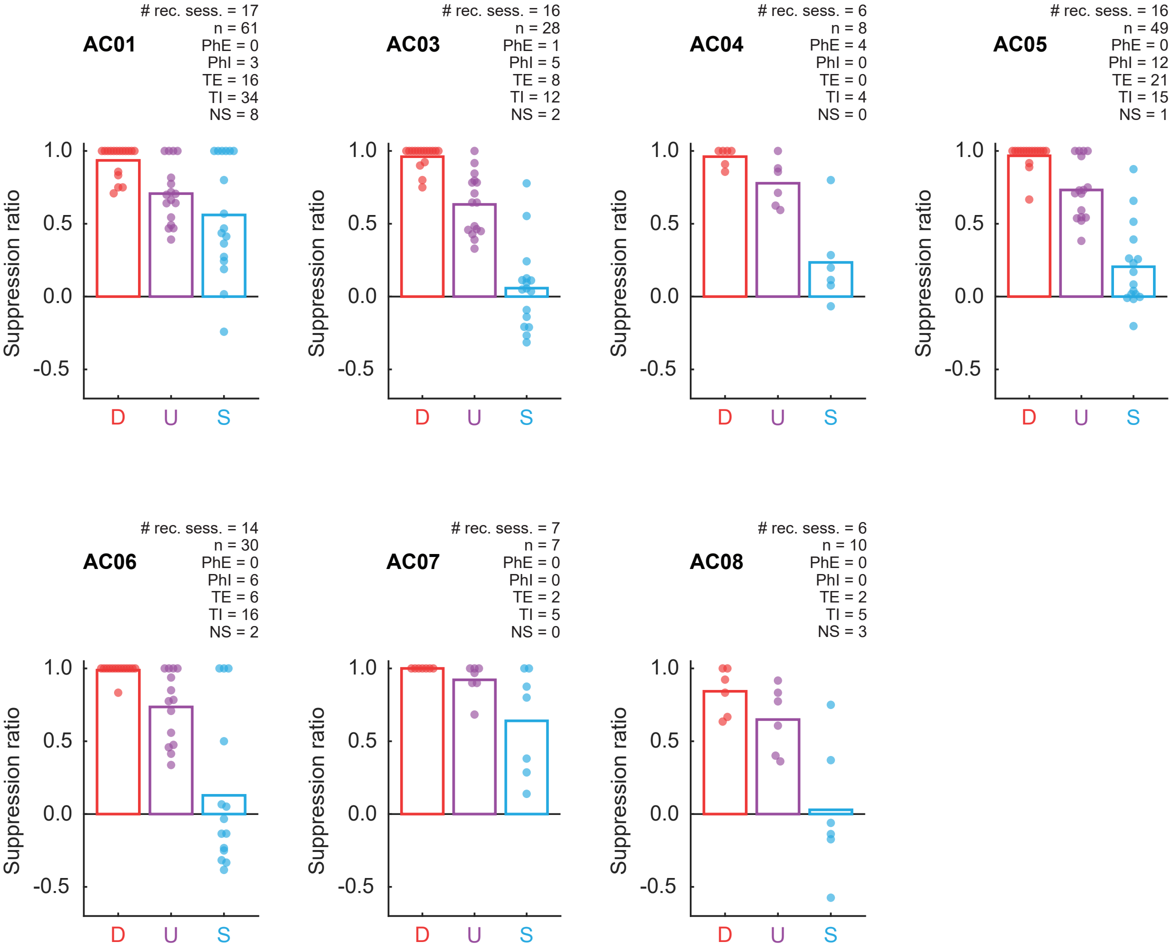


**Figure 2-1. Individual fear discrimination and recording summary.** Mean (bar) and individual session (data points) suppression ratio for each cue (D, danger, red; U, uncertainty, purple; S, safety, blue) is shown for each individual for all recording sessions with cue-responsive neurons. Animal identity is shown in the top left. For each individual, the number of recording sessions with cue-responsive neurons, the number of cue-responsive neurons, and the number of neurons in each cluster: PhE (Phasic Excited), PhI (Phasic Inhibited), TE (Tonic Excited), TI (Tonic Inhibited), and NS (Non-Selective) are provided.

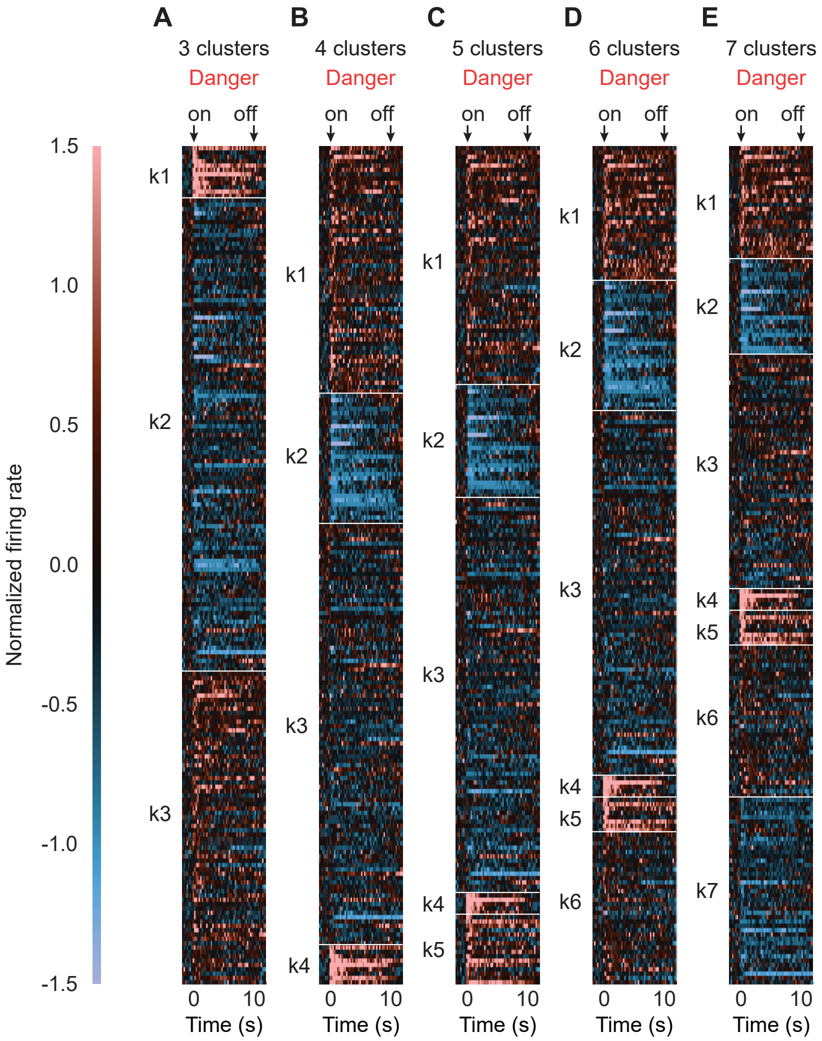


**Figure 2-1. Alternative clustering.** Heat plot showing mean normalized firing rate to danger (red) for each cue-responsive neuron (n = 193), from 2 s prior to cue onset to 2 s following cue offset. A normalized firing rate of zero is indicated by the color black. Firing increases in light red and firing decreases in light blue. Results for selecting (**A**) 3 clusters, (**B**) 4 clusters, (**C**) 5 clusters, (**D**) 6 clusters, and (**E**) 7 clusters. Cue onset (on) and offset (off) are indicated by black arrows.

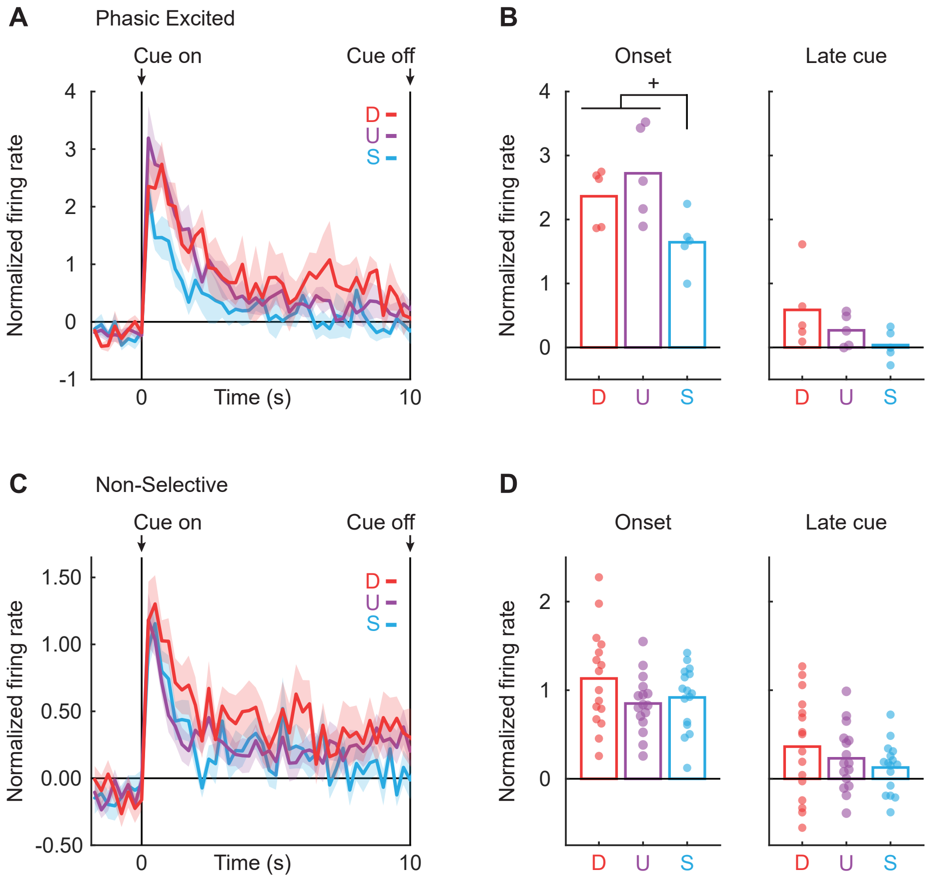


**Figure 2-2. Phasic Excited NAcc neurons are threat-responsive.** (**A**) Mean normalized firing rate to danger (D, red), uncertainty (U, purple), and safety (S, blue) is shown from 2 s prior to cue onset to cue offset for the Phasic Excited neurons (n = 5). Cue onset and offset are indicated by vertical black lines. (**B**) Mean (bar) and individual (data points), normalized firing rate for Phasic Excited neurons during the first 1 s cue interval (onset, left) and the last 5 s cue interval (late cue, right) are shown for each cue (D, danger; U, uncertainty; S, safety). Colors maintained from A. (**C**-**D**) Identical graphs made for Non-Selective neurons (n = 16), as in A and B. $^{+}$95% bootstrap confidence interval for differential cue firing does not contain zero.

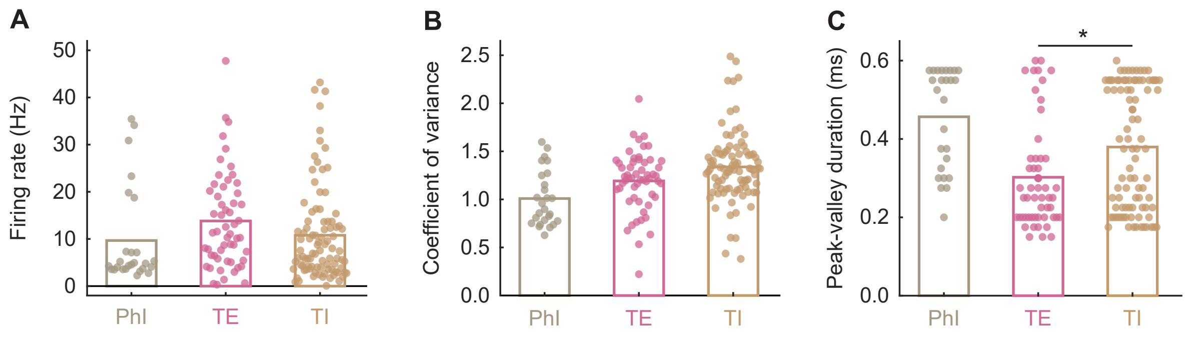


**Figure 2-3. Firing and waveform characteristics of cue-responsive neurons.** Mean (bar) and individual (data points) (**A**) firing rate, (**B**) coefficient of variance, and waveform peak-valley duration during a 10 s baseline period just prior to cue onset for Phasic Inhibited (PhI, n = 26, grey), Tonic Excited (TE, n = 55, pink), and Tonic Inhibited (TI, n = 91, tan) neurons. *Independent samples t-test; $p$ = 0.003.
